# CycPeptMP: Enhancing Membrane Permeability Prediction of Cyclic Peptides with Multi-Level Molecular Features and Data Augmentation

**DOI:** 10.1101/2023.12.25.573282

**Authors:** Jianan Li, Keisuke Yanagisawa, Yutaka Akiyama

## Abstract

Cyclic peptides are versatile therapeutic agents with many excellent properties, such as high binding affinity, minimal toxicity, and the potential to engage challenging protein targets. However, the pharmaceutical utilities of cyclic peptides are limited by their low membrane permeability—an essential indicator of oral bioavailability and intracellular targeting. Current machine learning-based models of cyclic peptide permeability show variable performance due to the limitations of experimental data. Furthermore, these methods use features derived from the whole molecule which are used to predict small molecules and ignore the unique structural properties of cyclic peptides. This study presents CycPeptMP: an accurate and efficient method for predicting the membrane permeability of cyclic peptides. We designed features for cyclic peptides at the atom-, monomer-, and peptide-levels, and seamlessly integrated these into a fusion model using state-of-the-art deep learning technology. Using the latest data, we applied various data augmentation techniques to enhance model training efficiency. The fusion model exhibited excellent prediction performance, with root mean squared error of 0.503 and correlation coefficient of 0.883. Ablation studies demonstrated that all feature levels were essential for predicting membrane permeability and confirmed the effectiveness of augmentation to improve prediction accuracy. A comparison with a molecular dynamics-based method showed that CycPeptMP accurately predicted the peptide permeability, which is otherwise difficult to predict using simulations.

## Introduction

Cyclic peptides are increasingly valued in pharmaceutical research due to their high binding affinity, target selectivity, and ability to inhibit intracellular protein–protein interactions (1, 2). The unique structural features of macrocyclic peptides stem from their restricted conformational flexibility and local secondary structure motifs, which allow for bioactive conformations with remarkable potency and selectivity (3). In the past century, cyclic peptide drugs were predominantly sourced from natural products, including antimicrobial agents and human peptide hormones. Recent advances in synthesis and screening systems have led to break-throughs in cyclic peptide drug discovery. Novel screening and cyclization strategies have greatly promoted cyclic peptide drug development in the past two decades (4, 5). For example, the random nonstandard peptides integrated discovery (RaPID) system designs cyclic peptides from a diverse library including non-natural amino acids, enabling the synthesis and rapid selection of potent binders for therapeutic targets (6). More than 40 cyclic peptide drugs are currently in clinical use, and the FDA has approved approximately one macrocyclic peptide drug annually for the past 20 years (5, 7). Despite their pharmacological potential, cyclic peptides often exhibit poor membrane permeability, severely limiting their biological applications and development of orally available drugs (8). While the mechanism of membrane permeation by cyclic peptides remains unclear, cyclic peptides with “closed”-conformation in hydrophobic environments demonstrate enhanced permeability (9, 10). The “closed” conformation conceals polar groups through intramolecular hydrogen bonds and lipophilic side chains, contributing to their increased permeation efficiency. Drawing inspiration from the structure of Cyclosporin A, a naturally occurring N-methylated macrocyclic peptide with high permeability, the shielding of the exposed hydrogen bond donor (-NH) through N-methylation has been widely applied to enhance membrane permeability (11, 12). Various strategies have emerged for improving membrane permeability, such as conformational control (2, 13), amide-to-ester substitution (14), amide-to-thioamide substitution (3), and side-chain modifications (15). However, these strategies do not improve membrane permeability across all cyclic peptides.

In the early stages of drug development, it is important to select candidate compounds with high membrane permeability. Because it is costly to randomly measure numerous peptides using biochemical assays, a rapid computational method for predicting membrane permeability is eagerly anticipated. Computational approaches for predicting the permeability of cyclic peptides have primarily been based on molecular dynamics (MD) simulation (16–19). Markov state models, among others, have been used to analyze simulation data and elucidate cyclic peptide behavior (20), which is crucial for understanding membrane permeation mechanisms and optimizing structures for enhancing membrane permeability. However, the computational cost of MD-based methods is a major limitation. In contrast to MD-based methods, several physicochemical or machine learning models have been developed, offering more rapid prediction capabilities (21–25). The descriptors for hydrophobicity, such as the octanol-water partition coefficient (LogP), are generally the most important features for prediction. However, all of these models were established using limited data sets (10– 250 cyclic peptides) and lack a sufficient degree of generalization performance. Furthermore, these methods directly apply whole-molecule features used for predicting the membrane permeability of small molecules, ignoring the unique structural characteristics of cyclic peptides, such as sequence information and circularity.

Unlike conventional physicochemical and machine learning approaches, deep learning (DL) models offer an architectural design tailored to peptide characteristics (26) and can automatically extract more complex structural features than small-molecule compounds from datasets. DL-based small molecule property prediction methods based on graph neural networks (GNNs) and transformers have become a major research area. By representing molecule atoms as nodes and bonds as edges, the GNN-based method can effectively capture molecular structural information and integrate the topological structure of molecules with complex atomic features. Nonetheless, most existing approaches, such as GCN (27, 28), GAT (29, 30), and MPNN (31), have intrinsic limitations, such as a poor ability to process global information and high risk of over-smoothing. In contrast, following the successful experience in natural language processing, many transformer-based methods have been proposed that treat SMILES as strings (32–34). Since these methods lack structural information, several methods have been developed using molecular graph representations as input, which can encode more complex atomic and bond information than strings (35–38). However, cyclic peptide permeability datasets are limited and diverse, and contain discrepancies and errors stemming from the use of various assay systems. To address these limitations, we constructed CycPeptMPDB (39): a comprehensive database of cyclic peptide membrane permeability. CycPeptMPDB comprises information on a total of 7334 cyclic peptides, including structures and experimentally measured membrane permeabilities, from 45 published studies and 2 patents from pharmaceutical companies, representing the first platform for developing DL-based prediction methods. Interestingly, over 99.6% of cyclic peptides include non-natural amino acids, which suggests that their creators aimed to enhance permeability through chemical modifications such as N-methylation, or by deliberately incorporating non-natural building blocks in their design.

Cyclic peptides are characterized by complicated conformational dynamics, where even a minor alteration in a single residue can lead to substantial changes in their membrane permeability (40). Therefore, many publications in CycPeptMPDB have focused on measuring changes in membrane permeability while varying only a few residues and maintaining a largely constant sequence. In the prediction of small molecule properties (38) and peptide–protein binding (41), the combined use of multi-scale molecular features can improve prediction accuracy. This study proposes CycPeptMP: a membrane permeability prediction model for cyclic peptides that effectively integrates multi-level features with state-of-the-art DL techniques. We engineered features at the peptide, monomer (substructures, such as residues), and atom levels to concurrently capture both the local sequence variations and global conformational changes in cyclic peptides. Additionally, to enhance the training efficiency of our model for complex cyclic peptides, we employed data augmentation methods at the three scales.

## Materials and Methods

### Experimental dataset

We used the structure and membrane permeability (LogP_exp_) of peptides in CycPeptMPDB. CycPeptMPDB contains membrane permeability data based on the parallel artificial membrane permeability (PAMPA), Caco-2, Madin–Darby canine kidney (MDCK), and Ralph Russ canine kidney (RRCK) assays. We selected PAMPA entries with the largest number of data points. If the same peptide was measured in multiple publications, we used the value recorded in the latest publication. Consequently, 6889 peptides were selected, covering an extremely wide range of molecular weights, from 342.44 to 1777.74. Considering that the lower limit of LogP_exp_ in CycPeptMPDB was −10 (1 *×* 10^−10^cm*/*s), but the detection limit in most publications was − 8 (1 *×* 10^−8^cm*/*s), we rounded the permeabilities of 314 peptides with values lower than − 8 to − 8. Similarly, the permeability of one peptide with a value higher than − 4 was rounded to − 4.

The validation and test sets were extracted from the overall data for model evaluation. First, the Kennard-Stone (KS) algorithm was employed to extract 5% of all data (344 peptides) as the test set, which should uniformly cover the multi-dimensional space (42). We generated 2048-bit Morgan fingerprints (Morgan FP, radius: 2) and selected the test set such that the Euclidean distance between each data point was maximized by the KS algorithm. From the remaining data, we randomly extracted 5% three times for the validation set (344 peptides), with no overlap between datasets. The membrane permeability and molecular weight distributions for each set are shown in Figure 1. The average root mean squared error (RMSE) and correlation coefficient (R) from three repeated runs were used as evaluation metrics.

**Fig. 1.**
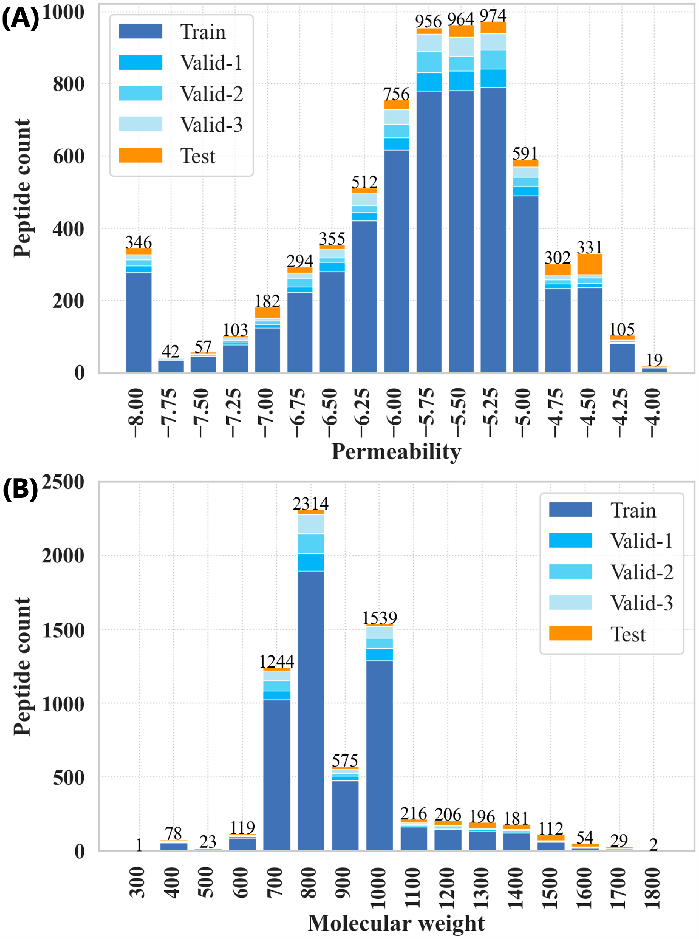
Experimental data distribution. (A) Membrane permeability. (B) Molecular weight. Valid–1 is the dataset used for the first-time evaluation of the validation set; the corresponding training data sets are Train, Valid–2, and Valid–3.

### Overview of CycPeptMP framework

Figure 2 shows the overall architecture of the CycPeptMP model. We designed three-level peptide representations and used each for three different sub-models to extract the atom-, monomer-, and peptide-level molecular representations. First, the input peptide was divided into monomers, and respective 3D conformations were generated from the peptide and monomers. Next, atom- and peptide-level features were extracted from the peptide conformation and used as input for the atom and peptide models, respectively. Monomer-level features extracted from the monomer conformation were used as input for the monomer model. Finally, the three-level latent feature vectors extracted using the three sub-models were combined to derive the membrane permeability prediction value.

**Fig. 2.**
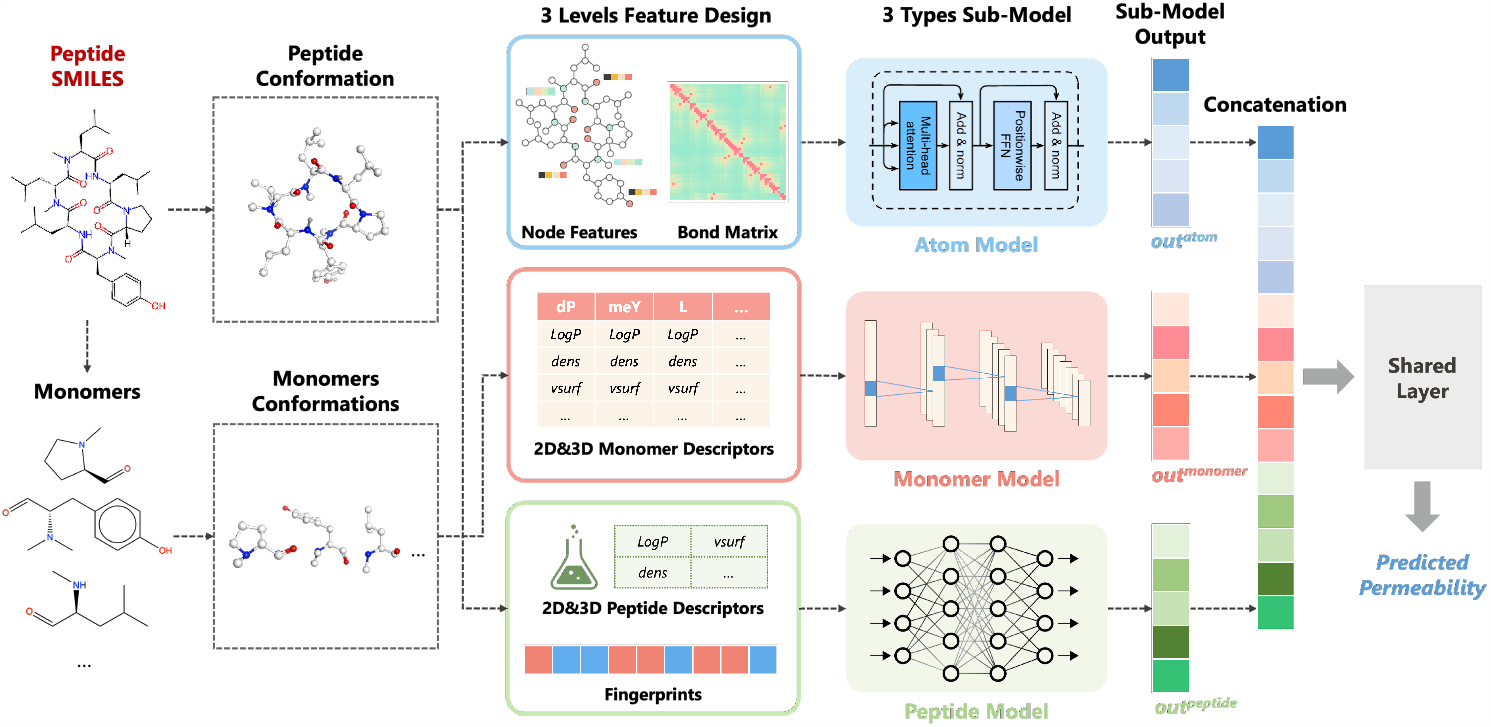
Overall framework of CycPeptMP model. The model incorporates the Transformer-based atom, convolutional neural network (CNN)-based monomer, and MLP-based peptide sub-models. The three-level expression vectors extracted by the three sub-models are concatenated and passed through a shared layer to derive the final permeability prediction value.

### Division of the monomer

We designed monomer-level features to accurately capture the subtle structural variations of cyclic peptides. The definition of a monomer corresponds to that provided by CycPeptMPDB. Although the peptide and ester bonds on the side chain were cleaved in CycPeptMPDB, bonds existing anywhere other than the macrocycle were not subjected to division to completely express the properties of the local structure. Merely hydrolyzing the peptide bond could generate a new hydrogen bond donor, which could misrepresent the original physicochemical properties of the substructure. Hence, when generating the conformation and calculating the monomer descriptor, the cleaved amide group or O atom of the amide-to-ester substitution was methylated (addition of CH3), and the carboxyl group was converted to an aldehyde group (addition of H).

### Peptide and monomer descriptors

To design peptide- and monomer-level features, peptides and monomers were represented by 16 descriptors, including LogP and polar surface area (Table 1, the correlation matrix of the selected descriptors is shown in Supplementary Figure S1). First, the 3D conformations of peptides and monomers were generated using RDKit. The initial structures were generated using the ETKDG method and then optimized using the UFF force field. Then, the 2D and 3D descriptors were calculated; we used a single-conformation 3D structure in the calculation. The 16 descriptors were selected as follows: First, a total of 1857 descriptors (1689 2D and 168 3D descriptors) were calculated for both cyclic peptides (peptide descriptors) and each monomer (monomer descriptors) using MOE software (version 2019.01) (43), the Mordred package (version 1.2) (44), and RDKit package (version 2022.09.5) (45). For 2D and 3D peptide descriptors, we removed all descriptors with constant values among cyclic peptides. For descriptor pairs with an absolute correlation coefficient of 0.9 or more, we excluded the one with the lower correlation with permeability. Consequently, 407 (335 2D and 72 3D descriptors) peptide descriptors were selected. We constructed two random forest (RF) models for predicting permeability based on the 2D or 3D peptide descriptors. Seven 2D and nine 3D peptide descriptors were selected based on the feature importance determined by the RF models (Supplementary Figure S2). Finally, the same 16 monomer descriptors were selected, and both peptide and monomer descriptors were standardized based on the Z-score.

**Table 1.**
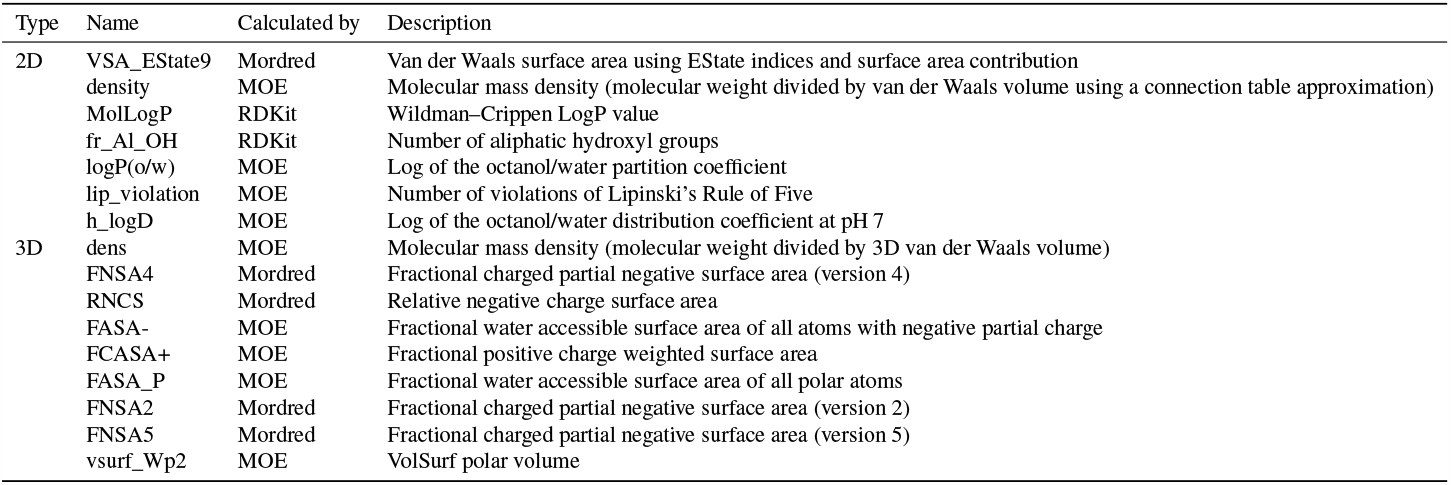
Selected descriptors, arranged in order of importance.

| Type | Name | Calculated by | Description |
| --- | --- | --- | --- |
| 2D | VSA_EState9 | Mordred | Van der Waals surface area using EState indices and surface area contribution |
|  | density | MOE | Molecular mass density (molecular weight divided by van der Waals volume using a connection table approximation) |
|  | MolLogP | RDKit | Wildman–Crippen LogP value |
|  | fr_Al_OH | RDKit | Number of aliphatic hydroxyl groups |
|  | logP(o/w) | MOE | Log of the octanol/water partition coefficient |
|  | lip_violation | MOE | Number of violations of Lipinski's Rule of Five |
|  | h_logD | MOE | Log of the octanol/water distribution coefficient at pH 7 |
| 3D | dens | MOE | Molecular mass density (molecular weight divided by 3D van der Waals volume) |
|  | FNSA4 | Mordred | Fractional charged partial negative surface area (version 4) |
|  | RNCS | Mordred | Relative negative charge surface area |
|  | FASA- | MOE | Fractional water accessible surface area of all atoms with negative partial charge |
|  | FCASA+ | MOE | Fractional positive charge weighted surface area |
|  | FASA_P | MOE | Fractional water accessible surface area of all polar atoms |
|  | FNSA2 | Mordred | Fractional charged partial negative surface area (version 2) |
|  | FNSA5 | Mordred | Fractional charged partial negative surface area (version 5) |
|  | vsurf_Wp2 | MOE | VolSurf polar volume |

### Atom model

We constructed a transformer-based atom model to capture the overall graph structure and 3D conformation of the peptide (Figure 3 (A)). We used node features (*Node*) and three types of bond weight matrices as inputs for the atom model (*Bond, Graph, Conf*). As shown in Table 2, heavy atoms were considered nodes and node features were represented as *Node*, bonded interaction weights were represented as 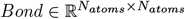, graphic pairwise distances were represented as 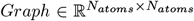, and 3D pairwise distances were represented as 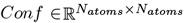.

**Table 2.** Input features of atom model.

| Type | Feature | Size | Description |
| --- | --- | --- | --- |
| Atom | Atom type | 8 | Type of atom by atomic number (one-hot) |
|  | Degree of atom | 5 | Number of bonds (one-hot) |
|  | Number of hydrogen | 5 | Number of bonded hydrogen atoms (one-hot) |
|  | Formal charge | 4 | Electrical charge (one-hot) |
|  | Hybridization | 3 | Type of atom hybridization (one-hot) |
|  | Chirality | 3 | Type of atom chirality (one-hot) |
|  | Is in a ring | 1 | Whether included in a ring structure |
|  | Is aromatic | 1 | Whether aromatic |
| Bond | Bond type (*Bond*) | 1 | Single(1.0), Double(2.0), Triple(3.0), Aromatic(1.5), Conjugated(1.4) and No-bond (0) |
|  | Graph distance (*Graph*) | 1 | Distance calculated from graph representation |
|  | 3D distance (*Conf*) | 1 | Euclidean distance (Å) calculated from 3D conformation |

**Fig. 3.**
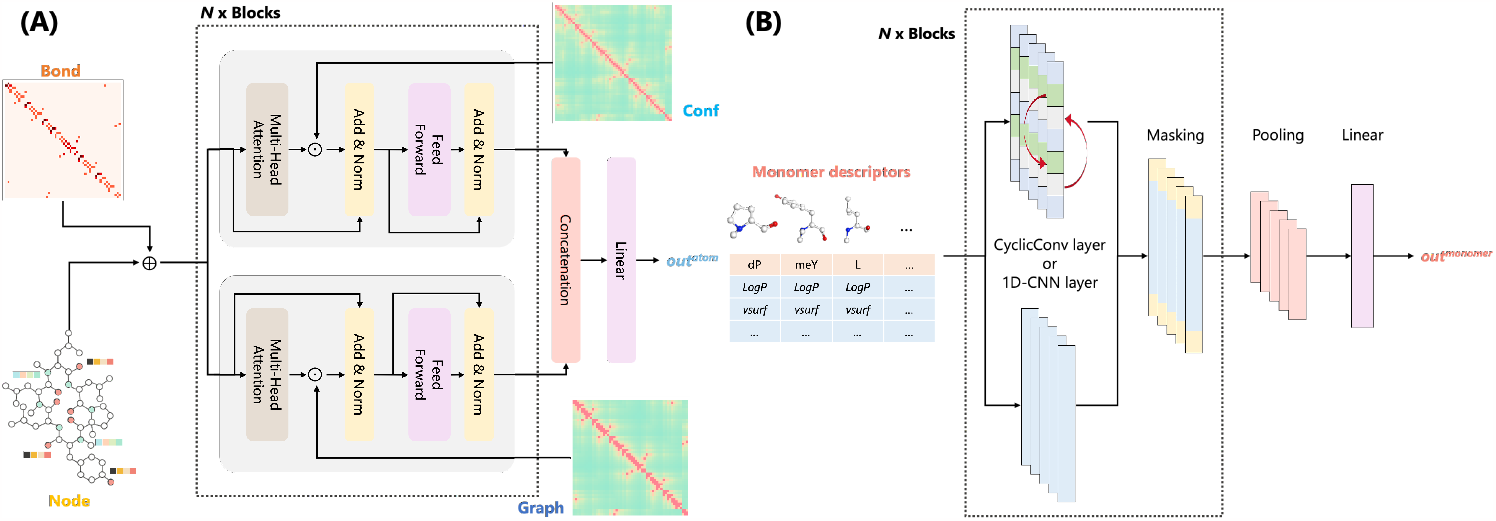
(A) Architecture of atom model. Node features and three types of bond matrices were used as input for the Transformer-based model. (B) Architecture of the monomer model. Monomer descriptors were used as input for the CNN-based model.

*Bond* recorded the molecule bond information and controlled message propagation between neighboring nodes by assigning weights to each bond type. According to the chemical bonding principle, bonds with more electron participation, such as unsaturated bonds, were assigned higher weights to enhance the exchange of information between atoms (37). To capture the local relationship between each node, embedded *Bond* was used for positional encoding and added to embedded *N ode* to serve as the input for the encoder block as follows:

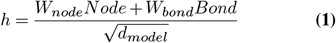

Where *d*_*model*_ is the attention dimension of the atom model, 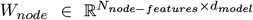 and 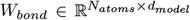 are trainable parameters, and 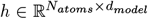 is the updated input for the encoder block.

Two types of distance matrices, *Graph* and *Conf*, the shortest pairwise graph distance and 3D Euclidean distance of each atom, were used to capture the overall structure and 3D conformation of the peptide. The distance maps were processed through an attenuation function to weaken distant interactions as follows:

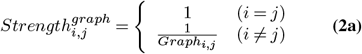

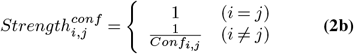

Here, we designed a structure-enhanced transformer encoder to learn the structural and 3D conformational information of peptides using focused attention. The encoder consisted of two parts, one using *Strength*^*graph*^ and another using *Strength*^*conf*^. This approach attenuates attention between less relevant pairs based on the distance, as shown in Equation (3) and Equation (4), providing a simplified approach to modeling complex molecular structures.

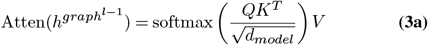

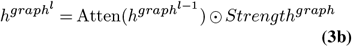

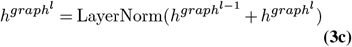

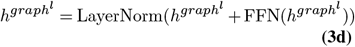

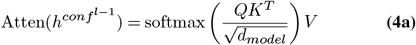

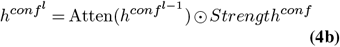

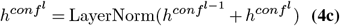

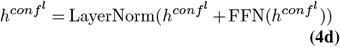

where 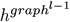 and 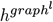 are the updated latent features of the *Graph* block (*l* − 1)-th and *l*-th layers, respectively, and 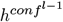 and 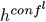 are the updated latent features of the *Conf* block (*l* − 1)-th and *l*-th layers, respectively.

Finally, the outputs 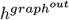 and 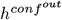 of the two blocks were weighted using the hyperparameter *λ*_*g*_ and the concatenated feature vector was used to derive the final output *out*^*atom*^ of the atom model as follows:

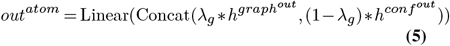

### Monomer model

To capture the partial structural information of peptides at the sequence level, we constructed a monomer model based on the 16 types of monomer descriptors (Figure 3 (B)). The CNN was used to learn partial structural features and sequence information. For the convolution layer, we used the general 1D-CNN layer or a CyclicConv layer (26) that considers peptide circularity. The use of the CyclicConv or 1D-CNN layer was determined by hyperparameter tuning. Finally, the monomer model derived the latent feature *out*^*monomer*^.

### Peptide model

The 16 peptide descriptors all expressed the physicochemical properties of the molecules. We also incorporated 2048-bit Morgan FP (1024-bit, radius: 2; 1024-bit, radius: 3) to express the substructural information for the peptide model. The descriptor and Morgan FP were each trained with different multilayer perceptrons (MLPs), and the latent feature vectors 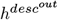 and 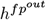 were concatenated and used to derive the final output *out*^*peptide*^ of the peptide model as follows:

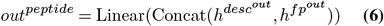

### Fusion model

As shown in Figure 2, the output latent feature vectors *out*^*atom*^, *out*^*monomer*^, and *out*^*peptide*^ of the three sub-models were concatenated to generate the final molecular feature vector, which was passed through a shared layer for the final permeability prediction *out*^*fusion*^. As the model becomes more complex, problems such as gradient disappearance may occur and input information may not be transmitted. Auxiliary loss is a learning technique in which additional losses are incurred to optimize the NN learning process. Directly propagating errors to the middle network layer can prevent gradient disappearance and improve embedding and learning efficiency (46). Hence, in addition to the main loss *L* calculated from the output of the fusion model, we designed the three sub-model losses *L*_*atom*_, *L*_*monomer*_, and *L*_*peptide*_ derived from the output of each sub-model, and layer losses 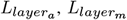, and 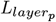 derived from the averaged outputs of the layers in each block (Transformer, CNN, and MLP) of the three sub-models. The loss function during training is presented in Equation (7):

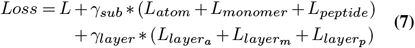

where the weight parameter *γ*_*sub*_ was set to 0.10 and *γ*_*layer*_ was set to 0.05. Only the output value *out*^*fusion*^ of the fusion model was used during inference. The hyperparameters of CycPeptMP were determined by 150 searches using Optuna software (47); the search range and results are shown in Supplementary Table S1.

### Data augmentation

Although the amount of available biological data has increased in recent years, experimental data remains limited compared to data for natural language processing and computer vision. For example, the Tox21 dataset, which deals with the toxicity classification of small molecules, only has about 8000 data points. This limitation in biological data has motivated the increased use of self-supervised learning approaches utilizing contrastive learning (48) and pre-training (32, 34). However, given a more limited availability compared to small molecules, these techniques remain challenging for cyclic peptides. To improve the learning efficiency of the model, we used three augmentation methods to generate 60 replicas based on the properties of SMILES, nature of cyclic peptide sequences, and the complexity of structural changes in cyclic peptides. First, the SMILES enumeration technique (49) was used to permute the atom order and generate input for the atom model with a different ordering. Next, considering the circularity of the cyclic peptide, the input of the monomer model was rearranged using sequence arrangement—the aligned monomer descriptors were translated and rotated as shown in Figure 4. Finally, 60 conformations per peptide/monomer were generated using RDKit to incorporate more diverse 3D information into the model. Cyclic peptide conformations were used to calculate the *Conf* matrix for the atom model and peptide descriptors for the peptide model, and monomer conformations were used to calculate the monomer descriptors for the monomer model. During training, each replica was given the same label and treated as independent data. During inference, the average of the 60 replicas was used as the final predicted value.

**Fig. 4.**
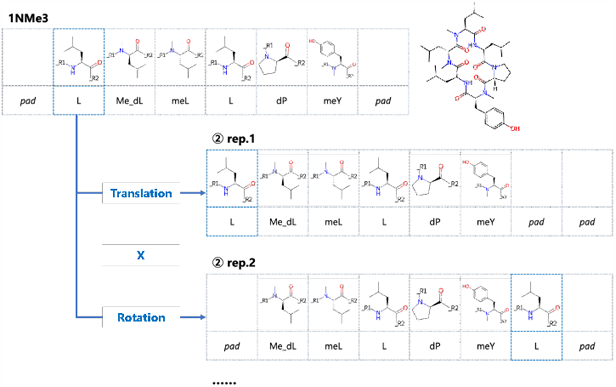
Sequence arrangement in the monomer model. The aligned monomer descriptors are translated and rotated based on the sequence information.

### Baseline methods

We validated the performance of CycPeptMP based on comparisons with seven baseline methods.

- Traditional baselines: To represent traditional cyclic peptide membrane permeability prediction methods, we constructed a RF model with 2048-bit Morgan FP, support vector machine (SVM) model with seven 2D peptide descriptors, and SVM model with 16 2D and 3D peptide descriptors. The hyperparameters of the RF and SVM models were determined by a grid search (Supplementary Table S2).
- Transformer-based methods: MAT (35) and SAT (36) were compared as state-of-the-art transformer-based methods for predicting small-molecule properties. MAT is a graph-based transformer that encodes the graph topology by adding weighted adjacency and 3D distance matrices to the self-attention. SAT is a structure-aware transformer that incorporates structural information into the original self-attention by extracting a subgraph representation rooted at each node before computing the attention.
- Multi-level feature methods: PharmHGT (38) designs features on the atom- and fragment-level and constructs a heterogeneous graph considering the correspondence between atoms and fragments for a transformer-based model. FinGAT (30) uses a GAT model to extract atom-level information and combines it with Morgan FP to capture the molecular structure from multiple perspectives.

The same validation set, test set, and random seeds of the DL-based model were used for all models to ensure a fair comparison.

## Results and Discussion

### Performance comparison

The prediction accuracy results for the test set are shown in Table 3, and the prediction accuracy results for the valid set are shown in Supplementary Table S3. CycPeptMP ranked first in all evaluation metrics (RMSE = 0.503, R = 0.883), reflecting a significant improvement in prediction performance over all existing methods. Considering the structural diversity of the test set, CycPeptMP showed good generalization performance and could learn the complex structures of cyclic peptides which is difficult to apply pre-training through augmentation. The RF model constructed based on Morgan FP showed good prediction performance and ranked third among all methods (RMSE = 0.616, R = 0.815). SVM with 2D peptide descriptors (RMSE = 0.653, R = 0.793) had lower prediction accuracy than the RF model, whereas SVM with 3D descriptors improved prediction accuracy, making it superior to the RF model for the test set (RMSE = 0.594, R = 0.833). Cyclic peptide membrane permeation by passive diffusion has a negative correlation with molecule size. SVM could predict permeability to some extent by using lipophilicity descriptors, such as LogP, which are largely dependent on molecular weight. CycPeptMP effectively combined Morgan FP and 16 2D and 3D peptide descriptors as peptide-level information to comprehensively characterize peptide structures from both a topological and physicochemical perspective, leading to an improvement in prediction capabilities. Neither graph representation transformer-based MAT (RMSE = 0.978, R = 0.408) nor SAT (RMSE = 1.071, R = 0.015) could predict membrane permeability. Although MAT and SAT are state-of-the-art methods for predicting small-molecule properties, they could not effectively learn the structures of more complex cyclic peptides. In addition to atom-level information, PharmHGT with fragment MACCS Keys (RMSE = 0.698, R = 0.771) and FinGAT with molecular Morgan FP (RMSE = 0.657, R = 0.788) had significantly improved prediction accuracies compared to MAT and SAT. FinGAT showed the same level of accuracy as the RF model and 2D SVM. These results revealed that designing features from various perspectives may be a key to successfully predicting the membrane permeability of cyclic peptides. These findings indicate that CycPeptMP effectively employs three levels of features to capture a wide range of structural information, from the smallest atomic detail to the broader peptide-level conformation.

**Table 3.** Performance comparison between CycPeptMP and seven baseline methods using the test set. The metrics are the averaged values of three repeated runs, and the best result for each metric is indicated in bold.

| Evaluation Metrics | RF | SVM (2D) | SVM (2D3D) | MAT | SAT | PharmHGT | FinGAT | CycPeptMP |
| --- | --- | --- | --- | --- | --- | --- | --- | --- |
| RMSE | 0.616 | 0.653 | 0.594 | 0.978 | 1.071 | 0.698 | 0.657 | **0.503** |
| R | 0.815 | 0.793 | 0.833 | 0.408 | 0.015 | 0.771 | 0.788 | **0.883** |

Peptides with an experimental value of − 8, the lower limit, could not be predicted by CycPeptMP (Figure 5, results for the three validation sets are shown in Supplementary Figure S3). Compared to the RMSE of all data (0.503), the RMSE decreased significantly for data excluding peptides with the experimental value of 8 (0.468). Due to the detection limits of the different assay environments, we rounded the data from 8 to 10 to 8, though this may not be the only suitable approach.

**Fig. 5.**
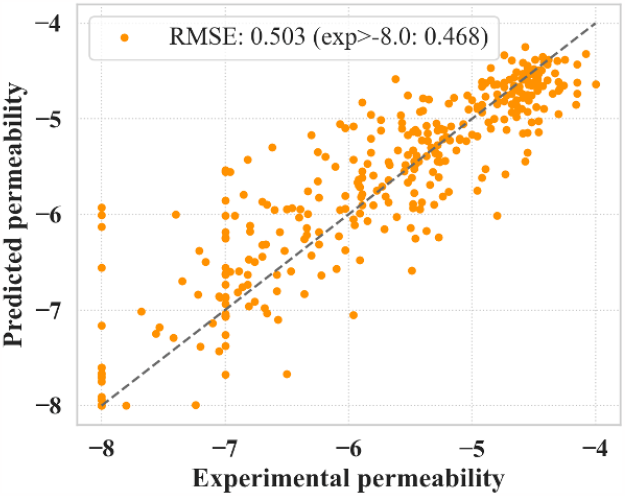
CycPeptMP prediction results for the test set. The predicted value of the test set is the average value of three runs.

### Ablation study of atom and monomer models

We conducted ablation studies on atom and monomer models with complex architectures (Figure 6, the results for the validation set are shown in Supplementary Figure S4 (A)). For the atom model, A is the original model and A–aug is the result without data augmentation. We measured the prediction accuracy when not using the *Bond* matrix (A–bond), using ordinary absolute positional encoding (50) instead of the *Bond* matrix (A–abpe), retaining only the *Conf* block (A–graph), or retaining only the graph block (A–conf). As shown in Figure 6, the prediction accuracy of the atom model was significantly improved by augmentation (A: 0.653, A– aug: 0.931). Regarding the architecture changes of the atom model, the original A showed the highest prediction accuracy, and any element deletion decreased the prediction accuracy. It was better to capture the relationship between atoms using *Bond* (A: 0.653) than absolute positional encoding (A–abpe: 0.678), and the impact of removing *Graph* block (A–graph: 0.67) was greater than that of removing *Conf* block (A–conf: 0.659).

**Fig. 6.**
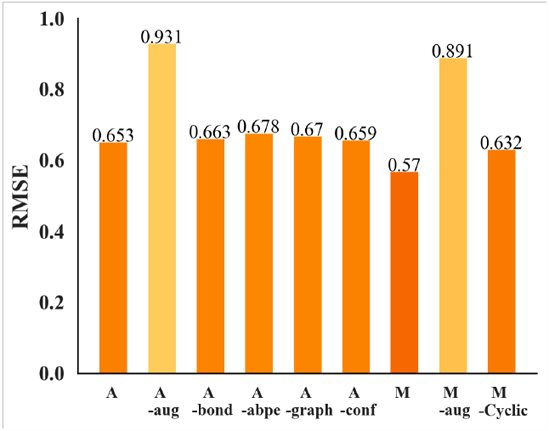
Ablation results (RMSE) for the atom and monomer models using the test set.

For the monomer model, M is the original model and M–aug is the result without data augmentation. We also measured the change in accuracy when replacing the general 1D-CNN layers with CyclicConv layers (M–Cyclic). Similar to the atom model results, the prediction accuracy of the monomer model was significantly improved by augmentation (M: 0.57, M–aug: 0.891). These results showed that SMILES enumeration for the atom model and sequence arrangement for the monomer model effectively improved learning efficiency, and that the augmentation technique is essential for learning the complex structure of cyclic peptides. Additionally, the 1D-CNN layer (M: 0.57) was superior to the CyclicConv layer (M–Cyclic: 0.632), consistent with previous findings (26).

### Ablation study of fusion model

The ablation study for the fusion model measured the influences of the number of replicas generated by augmentation and changes in architecture. Figure 7 (A) shows the accuracy based on 1 (no augmentation), 5, 10, 20, 30, 40, 50, and 60 (CycPeptMP) replicas per peptide. Even with five replicas (F–5: 0.544), we observed a significant improvement in prediction accuracy compared to that without augmentation (F–1: 0.61). However, over 20 replicas showed approximately the same prediction accuracy as the amount of training data increased, which may be due to the limitations of increasing diversity by mere reordering of inputs and lack of diversity in the generated conformations. In Figure 7 (B), F is the original CycPeptMP model; F–aux is that without auxiliary loss; F–atom, F–mono, and F–pep are those lacking the respective sub-models; F–3D is that not using all 3D information (*Conf* and 3D descriptors); and F– ensem is that where each sub-model is allowed to directly predict membrane permeability and the average ensemble of three predictions is taken. Auxiliary loss did not improve prediction accuracy (F–aux: 0.501). Furthermore, the prediction accuracy decreased when removing any of the three sub-models, indicating that the three levels of information is essential for predicting membrane permeability. The peptide model had the greatest influence (F–pep: 0.539), followed by the monomer (F–mono: 0.53) and atom model (F–atom: 0.511). The use of 3D information improved prediction accuracy (F–3D: 0.528), though not significantly so. Accuracy may be improved by generating conformations using a more rigorous method, such as MD simulations. Finally, the average prediction accuracy decreased further when using a sub-model ensemble (F–ensem: 0.529). This indicates that, rather than having each sub-model predict permeability directly, it was better to extract latent features.

**Fig. 7.**
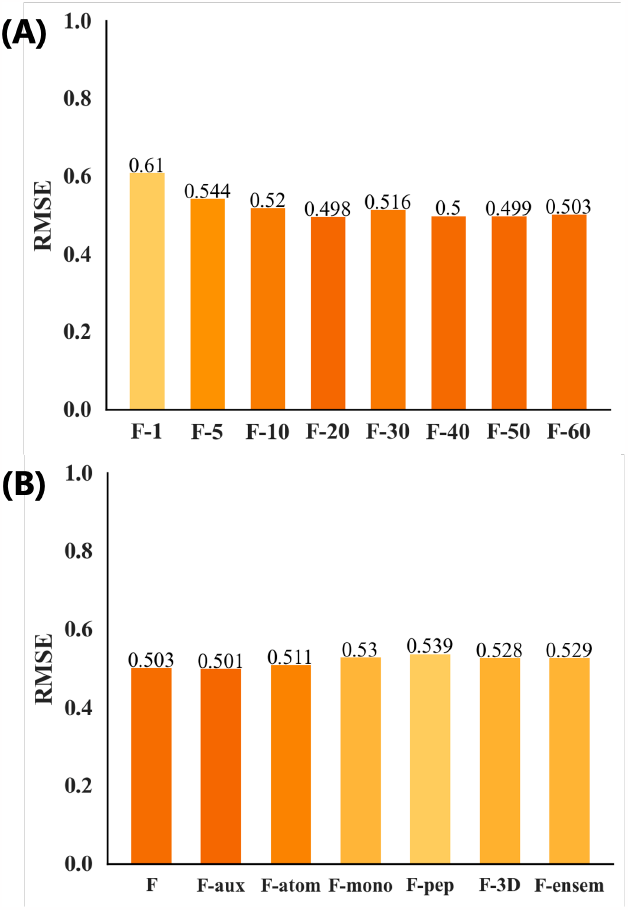
Ablation results (RMSE) for the fusion model using the test set. (A) Results for different numbers of input replicas. (B) Results for different architectures.

### Comparison with MD-based method

Cyclic peptides tend to exist in various conformations, resulting in slow conformational transitions relative to simulation time scales. Sugita et al., in the first large-scale prediction of cyclic peptide membrane permeability (18), used steered MD and replica-exchange umbrella sampling to accelerate sampling and simulated the membrane permeation process of 100 six-residue and 56 eight-residue peptides through a lipid bilayer. We compared their prediction results with the CycPeptMP results for 23 peptides (Supplementary Table S4) included in three validation sets (16 peptides) and the test set (7 peptides). While the MD-based method could not predict the membrane permeability of these 23 peptides (RMSE = 1.899), CycPeptMP was able to make accurate predictions for all of them (RMSE = 0.127) (Figure 8). Because hydrophobic cyclic peptides have insufficient solubility, diffuse slowly in the unstirred water layer, and are likely adsorbed to the membrane, Sugita et al. could not reproduce these behaviors using the inhomogeneous solubility-diffusion model (ISDM), which considers only direct membrane permeation processes (18). Thus, they reported a prediction accuracy (R) of only 0.21 for all 100 six-residue peptides; however, when 33 hydrophobic peptides (AlogP *>*= 4) were excluded, the accuracy increased to 0.54. A similar trend was seen among the 23 peptides compared in this study: by excluding six peptides with AlogP *>*= 4, the RMSE of the MD-based method increased from 1.899 to 1.219. Overall, CycPeptMP could accurately and quickly predict peptide permeability with far superior performance compared to MD-based methods, and thus represents a promising tool for cyclic peptide drug discovery.

**Fig. 8.**
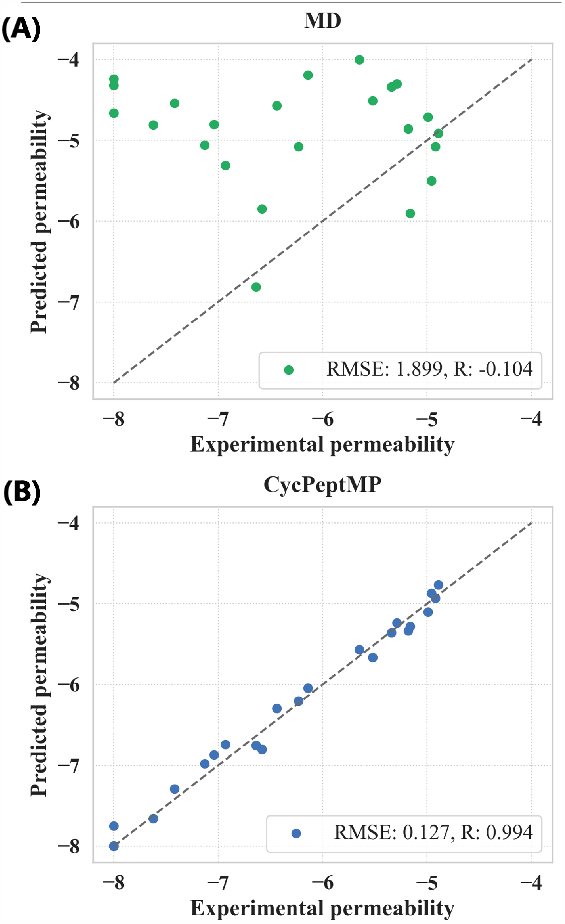
Prediction results of (A) MD-based method and (B) CycPeptMP.

## Conclusion

CycPeptMP represents a high-performance deep learning-based technique for predicting the membrane permeability of cyclic peptides. CycPeptMP incorporated atom-, monomer-, and peptide-level features, and improved training efficiency through three types of data augmentation techniques. CycPeptMP showed excellent prediction accuracy and generalization performance compared to existing methods. Moreover, we confirmed that CycPeptMP was able to accurately predict the permeability of peptides where MD-based methods failed with much lower computational costs. Thus, CycPeptMP shows great potential for advancing cyclic peptide drug discovery through rapid identification of high-permeability peptides, and can inform the development of more effective DL-based techniques in related fields. Future studies should focus on improving prediction performance by generating 3D conformations with a more rigorous method.

### Key Points

- This study presents CycPeptMP—a novel DL-based method for predicting the membrane permeability of cyclic peptides. CycPeptMP utilizes a multi-level feature design and data augmentation to simplify the characterization of complex peptide structures and improve model performance.
- CycPeptMP achieved state-of-the-art performance using a test set containing a variety of structures, which demonstrates the functionality gained by implementing the three-level feature-appropriate architectural design.
- CycPeptMP has excellent performance for peptides that are difficult to predict with MD-based methods, and can promote the efficacy of cyclic peptide drug discovery in many research directions, such as structure–activity relationship analysis and lead optimization.

## Supporting information

latex source

## ACKNOWLEDGEMENTS

This work was partially supported by KAKENHI (Grant no. 20K19917, no. 22H03684, no. 23KJ0891, and no. 23H03495) of the Japan Society for the Promotion of Science (JSPS), and Basis for Supporting Innovative Drug Discovery and Life Science Research (BINDS) (Grant no. JP23ama121026j0002) of the Japan Agency for Medical Research and Development (AMED).

