## Supplementary Data for "CycPeptMP: Enhancing Membrane Permeability Prediction of Cyclic Peptides with Multi-Level Molecular Features and Data Augmentation"

1 **Supplemental Figures**

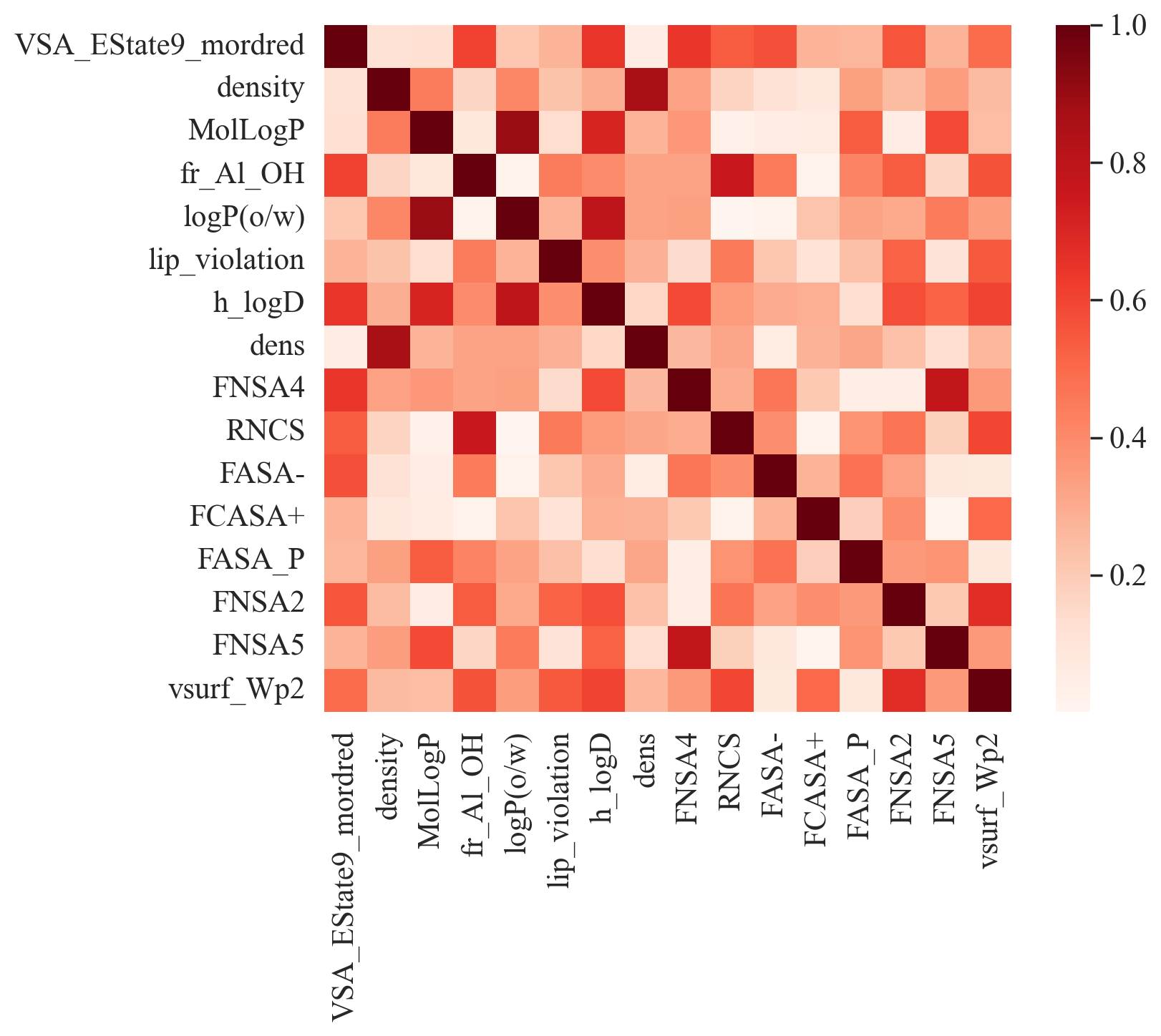
 **Figure** **S1**: Heat map of absolute correlation coefficient values for 16 peptide descriptors. The pair with the highest correlation is MolLogP and logP(o/w) (|R| = 0.898).

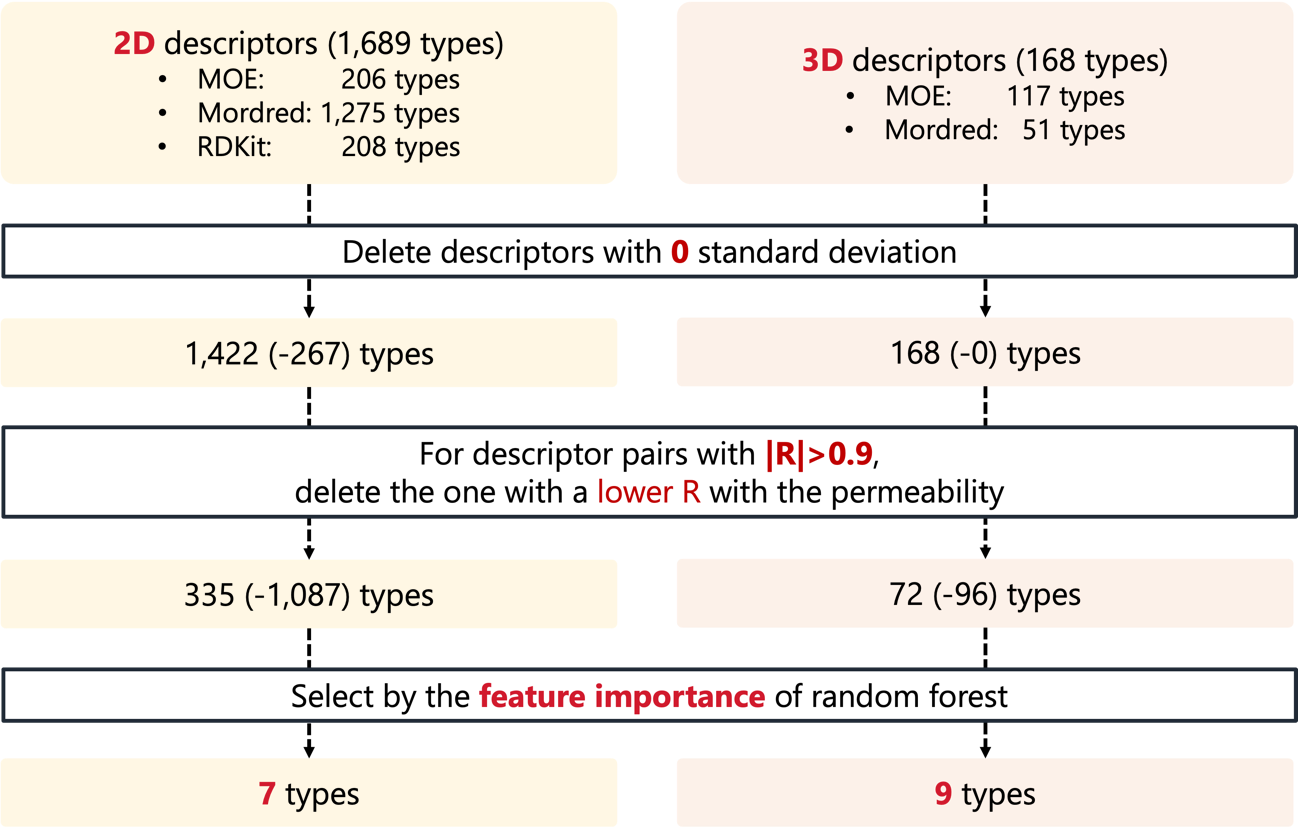

**Figure** **S2**: Preprocessing and selection of peptide descriptors. 206 2D descriptors and 117 3D descriptors were calculated by MOE [1], 1275 2D and 51 3D descriptors that could be calculated correctly were calculated by Mordred [2], and 208 2D descriptors were calculated by RDKit [3]. Finally, seven 2D peptide and nine 3D peptide descriptors were selected based on feature importance of a random forest model.

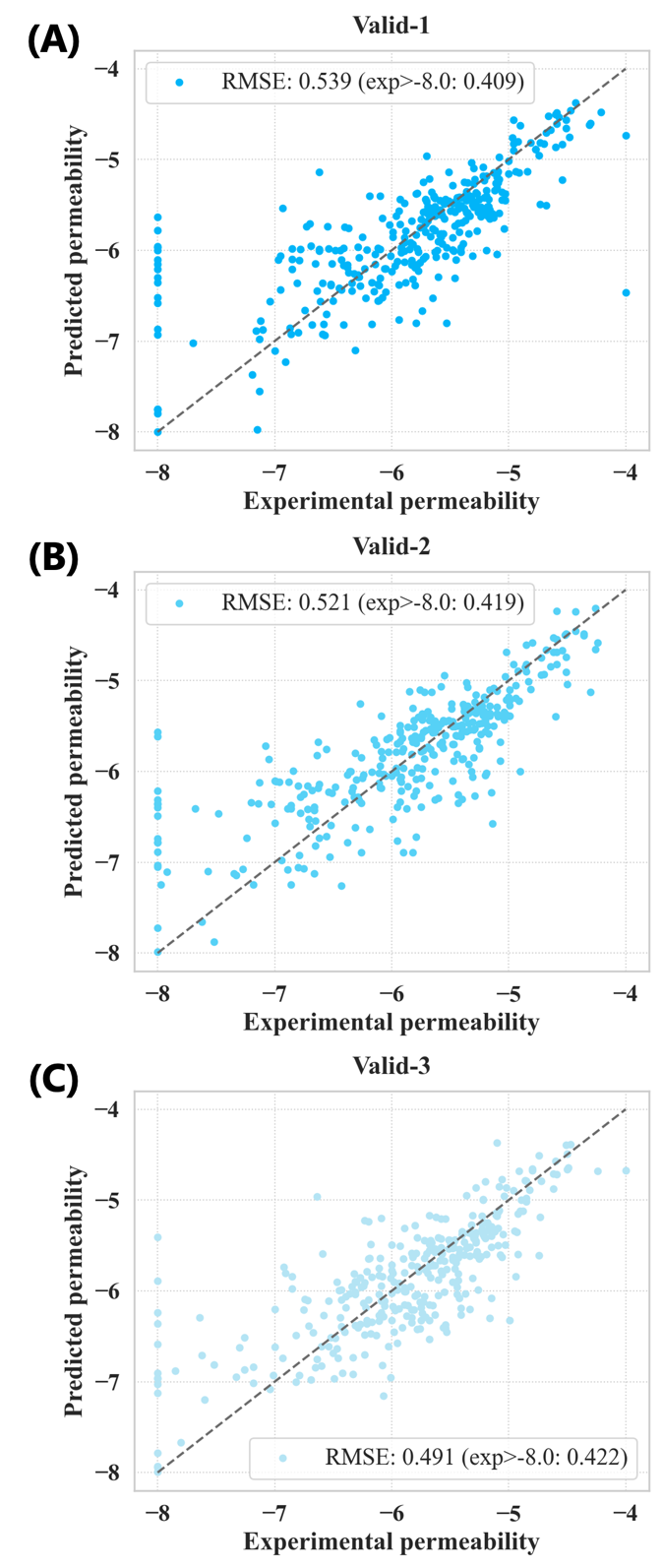

**Figure** **S3**: CycPeptMP prediction results of (A) 1st validation set (Valid-1), (B) 2nd validation set (Valid-2), and (C) 3rd validation set (Valid-3). Each figure shows the RMSE of all data and RMSE calculated after excluding data with an experimental value of –8, the lower limit, which could not be predicted with the three validation sets. The RMSE of all validation sets decreased significantly after excluding peptides with an experimental value of –8.

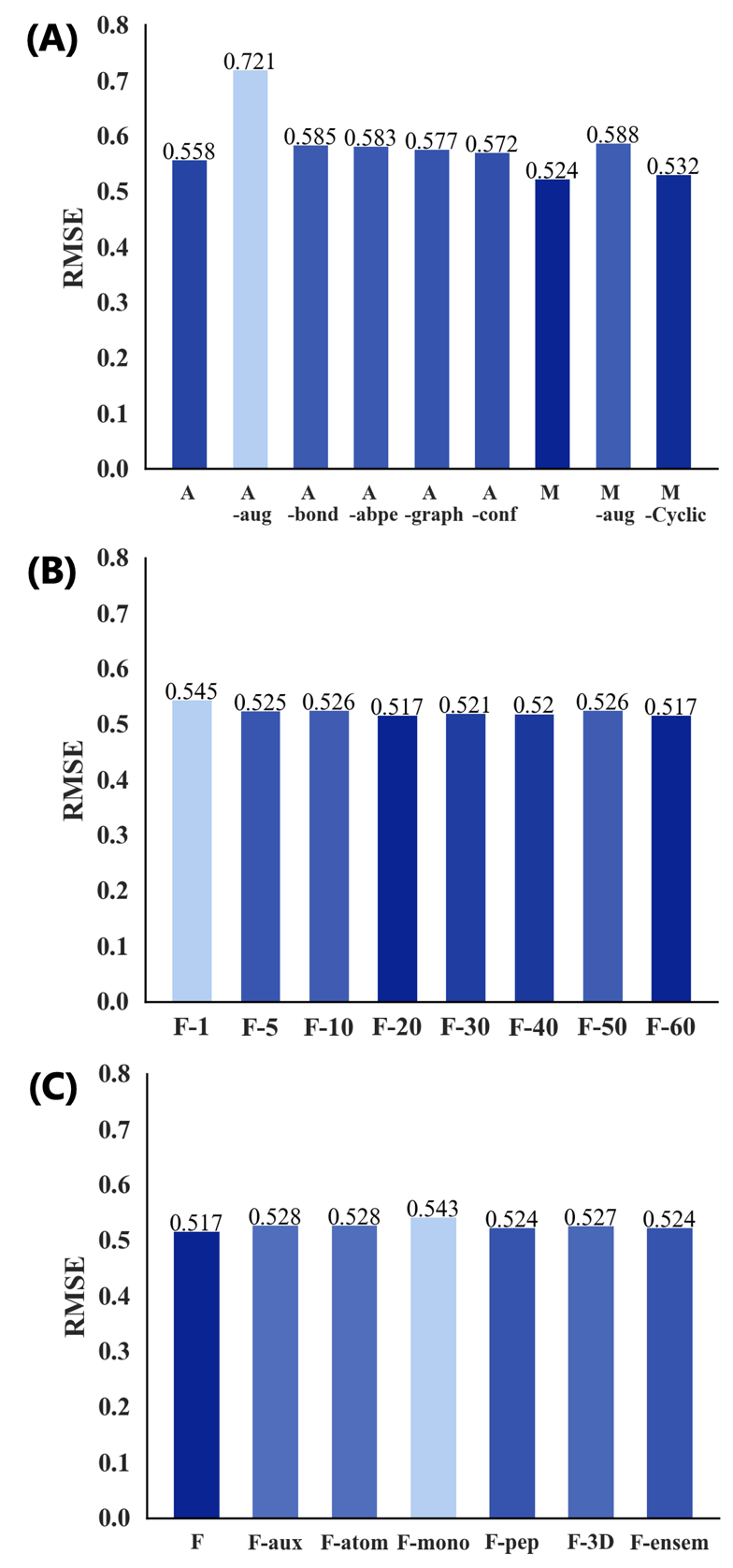

**Figure** **S4**: Ablation results (RMSE) for the atom and monomer models using the validation set (A), and ablation results for the fusion model (B, C). (A) Effects of changing the number of input replicas and architecture for the atom and monomer models. (B) Effects of changing the number of input replicas for the fusion model. (C) Effects of changing the architecture for the fusion model.

2 **Supplemental Tables**

**Table** **S1**: Results of hyperparameter search (150 searches using Optuna software [4]) for the CycPeptMP model.

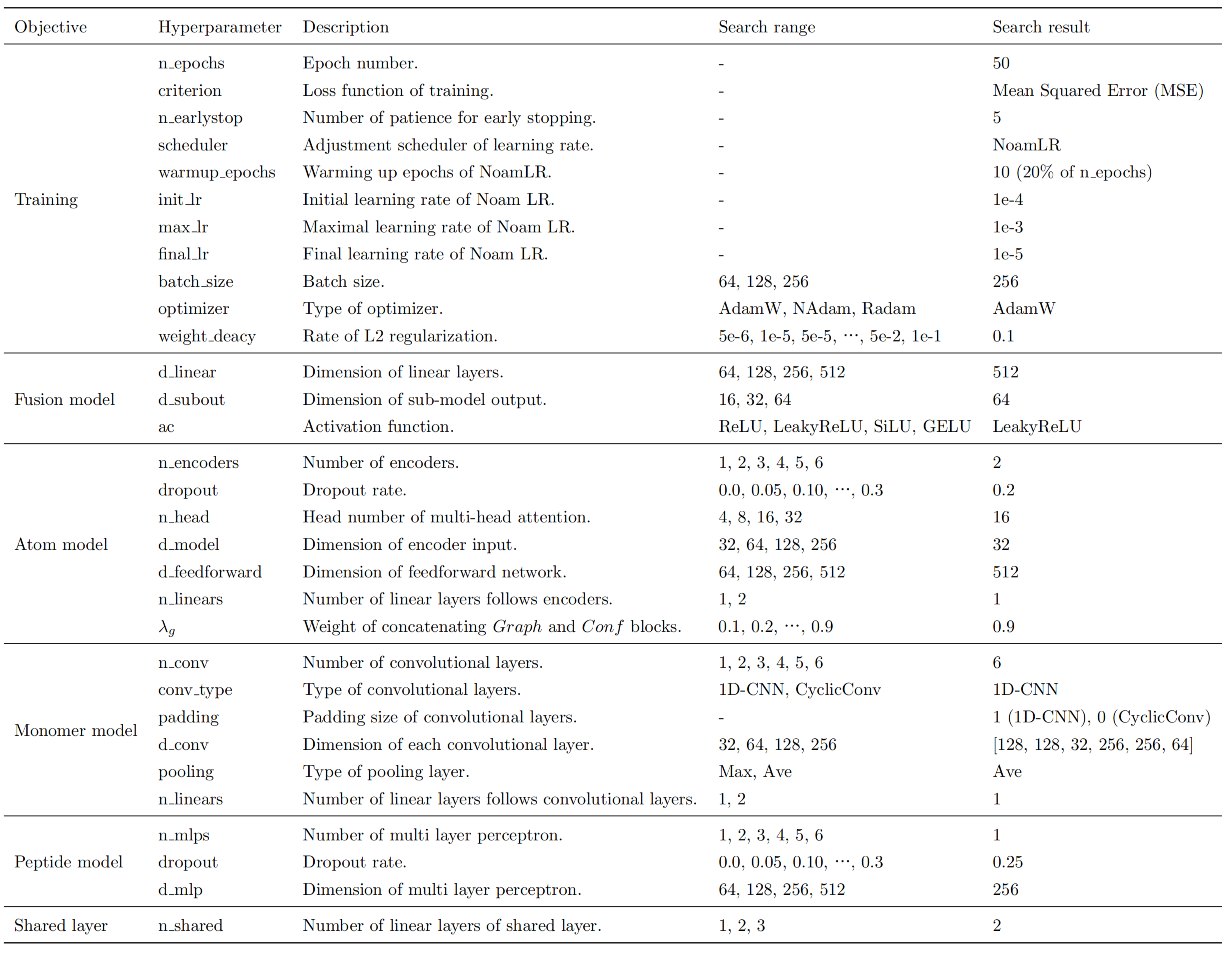

**Table** **S2**: Results of the hyperparameter search (grid search) for the RF and SVM models.

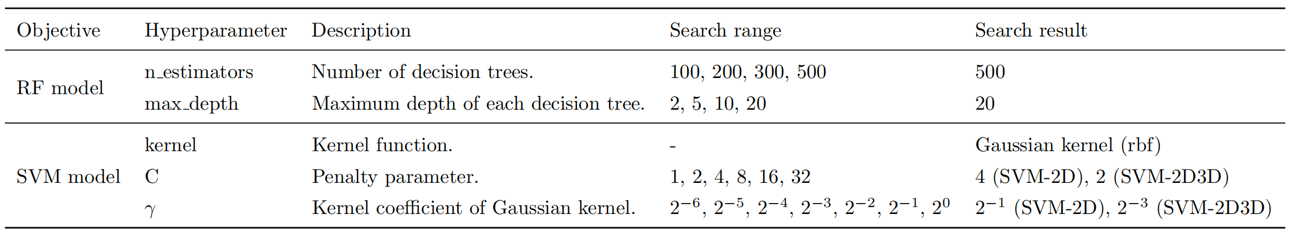

**Table** **S3**: Comparison of performance between CycPeptMP and seven baseline methods using the validation set; the metrics are averaged for three runs, and the best result for each metric is indicated in bold.

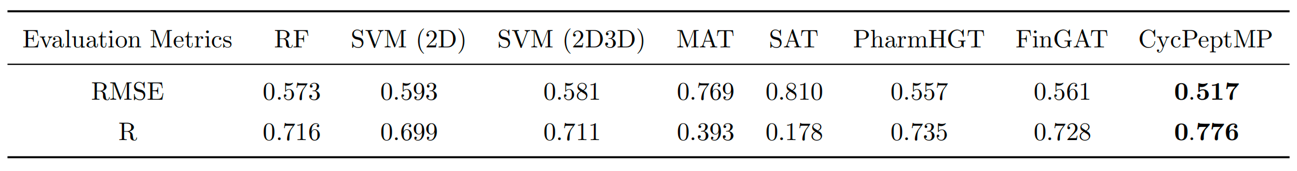

**Table** **S4**: Peptides used in comparison with MD-based method. The 23 peptides are included in the validation and test sets of this study. The AlogP and MD predicted value (${logP}_{ISMD\_mod}$) are reported by Sugita et al [5].

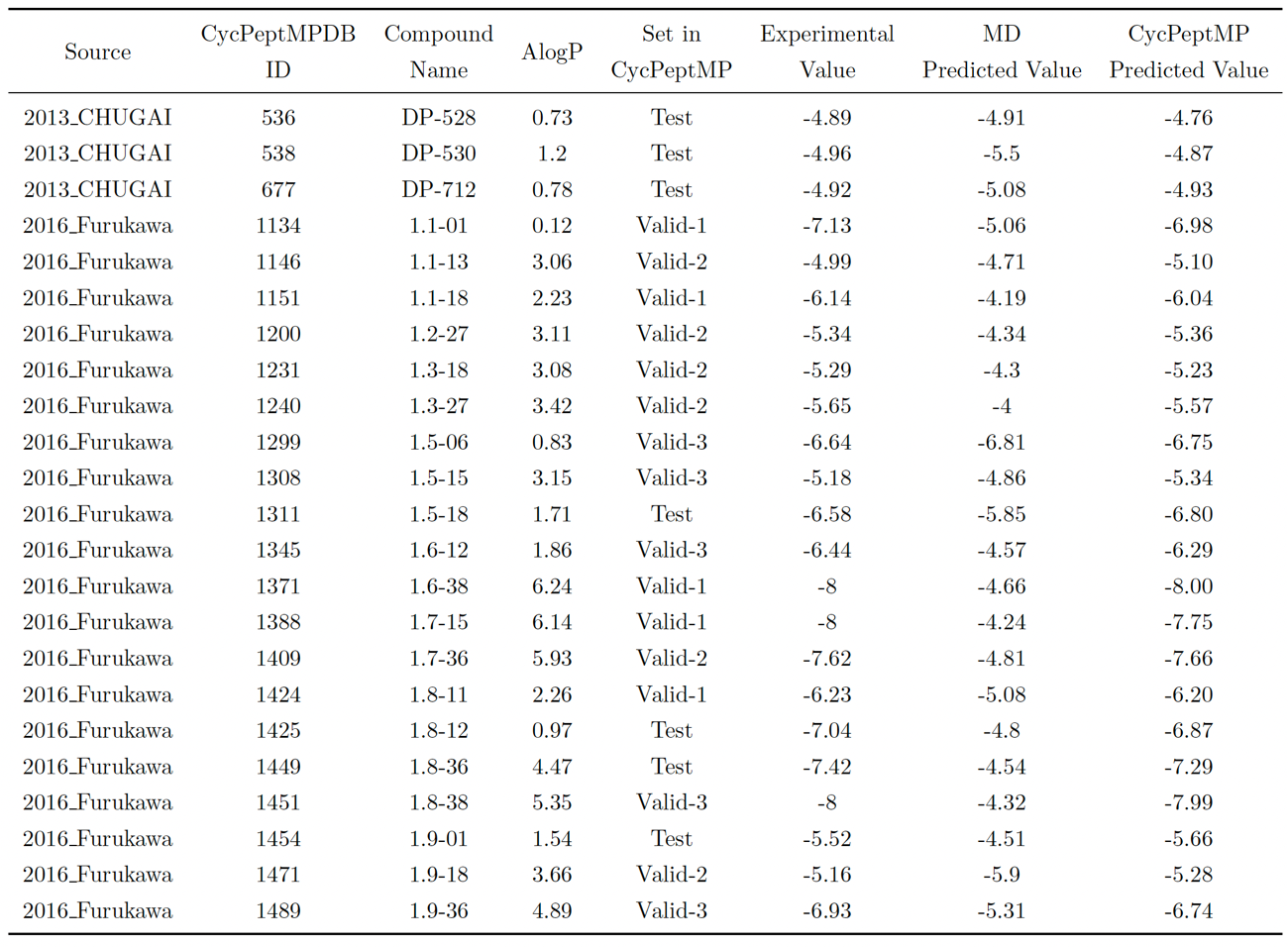
