## Supplementary material for "CycPeptMP: Enhancing Membrane Permeability Prediction of Cyclic Peptides with Multi-Level Molecular Features and Data Augmentation": latex source: Article.tex

\documentclass[times, twoside, watermark]{zHenriquesLab-StyleBioRxiv}
% \usepackage{blindtext}
\usepackage{booktabs}

\graphicspath{{Fig/}}


\def\SupFigDesccorr{Supplementary Figure~S1}
\def\SupFigDescpre{Supplementary Figure~S2}
\def\SupFigPlotValid{Supplementary Figure~S3}
\def\SupFigAblationValid{Supplementary Figure~S4}
\def\SupTabOptuna{Supplementary Table~S1}
\def\SupTabGrid{Supplementary Table~S2}
\def\SupTabResultsValid{Supplementary Table~S3}
\def\SupTabResultsMD{Supplementary Table~S4}


% Please give the surname of the lead author for the running footer
\leadauthor{Li} 

\begin{document}

\title{CycPeptMP: Enhancing Membrane Permeability Prediction of Cyclic Peptides with Multi-Level Molecular Features and Data Augmentation}
\shorttitle{CycPeptMP}

% Use letters for affiliations, numbers to show equal authorship (if applicable) and to indicate the corresponding author
\author[1]{Jianan Li}
\author[1, 2]{Keisuke Yanagisawa}
\author[1, 2, $\ast$]{Yutaka Akiyama}

\affil[1]{Department of Computer Science, School of Computing, Tokyo Institute of Technology, 2-12-1 Ookayama, Meguro-ku, 152-8550, Tokyo, Japan}
\affil[2]{Middle-Molecule IT-based Drug Discovery Laboratory (MIDL), Tokyo Institute of Technology, 2-12-1 Ookayama, Meguro-ku, 152-8550, Tokyo, Japan}


\maketitle

%TC:break Abstract
%the command above serves to have a word count for the abstract
\begin{abstract}
Cyclic peptides are versatile therapeutic agents with many excellent properties, such as high binding affinity, minimal toxicity, and the potential to engage challenging protein targets. 
However, the pharmaceutical utilities of cyclic peptides are limited by their low membrane permeability---an essential indicator of oral bioavailability and intracellular targeting. 
Current machine learning-based models of cyclic peptide permeability show variable performance due to the limitations of experimental data. 
Furthermore, these methods use features derived from the whole molecule which are used to predict small molecules and ignore the unique structural properties of cyclic peptides. 
This study presents CycPeptMP: an accurate and efficient method for predicting the membrane permeability of cyclic peptides.
We designed features for cyclic peptides at the atom-, monomer-, and peptide-levels, and seamlessly integrated these into a fusion model using state-of-the-art deep learning technology. 
Using the latest data, we applied various data augmentation techniques to enhance model training efficiency. 
The fusion model exhibited excellent prediction performance, with root mean squared error of $0.503$ and correlation coefficient of $0.883$. 
Ablation studies demonstrated that all feature levels were essential for predicting membrane permeability and confirmed the effectiveness of augmentation to improve prediction accuracy. 
A comparison with a molecular dynamics-based method showed that CycPeptMP accurately predicted the peptide permeability, which is otherwise difficult to predict using simulations.
\end {abstract}
%TC:break main
%the command above serves to have a word count for the abstract


\begin{keywords}
cyclic peptide | data augmentation | deep learning | feature design | membrane permeability
\end{keywords}


% \begin{corrauthor}
% %\texttt{r.henriques{@}ucl.ac.uk}
% akiyama\at c.titech.ac.jp
% \end{corrauthor}


%%%%%%%%%%%%%%%%%%%%%%%%%%%%

\input{Introduction}
\input{Methods}
\input{Results_and_Discussion}
\input{Conclusion}

%%%%%%%%%%%%%%%%%%%%%%%%%%%%


\begin{acknowledgements}
This work was partially supported by 
% 科研費
KAKENHI (Grant no. 20K19917, no. 22H03684, no. 23KJ0891, and no. 23H03495) of the Japan Society for the Promotion of Science (JSPS),
% % 学振
% Grant-in-Aid for JSPS Fellows (Grant no. 23KJ0891) of the JSPS,
% BINDSいるか？
and Basis for Supporting Innovative Drug Discovery and Life Science Research (BINDS) (Grant no. JP23ama121026j0002) of the Japan Agency for Medical Research and Development (AMED).
\end{acknowledgements}


\section*{Reference}
\bibliography{reference}

%% You can use these special %TC: tags to ignore certain parts of the text.
%TC:ignore
%the command above ignores this section for word count
\onecolumn
\newpage


% \section*{Word Counts}
% This section is \textit{not} included in the word count. 
% \subsection*{Notes on Nature Methods Brief Communication}
% \begin{itemize}
% \item Abstract: 3 sentences, 70 words.
% \item Main text: 3 pages, 2 figures, 1000-1500 words, more figures possible if under 3 pages
% \end{itemize}


% \subsection*{Statistics on word count}
% \detailtexcount
% \newpage


%%%%%%%%%%%%%%%%%%%%%%%%%%%%%
% Supplementary Information %
%%%%%%%%%%%%%%%%%%%%%%%%%%%%%


% \captionsetup*{format=largeformat}
% \section{Supplemental Figures} 

% \begin{figure*}[tb]
%     \centering
%     \includegraphics[width=\linewidth, natwidth=40.22cm, natheight=19.93cm]{Fig/SupFigDesccorr.png}
%     \caption{Heat map of absolute correlation coefficient values for $16$ peptide descriptors. The pair with the highest correlation is MolLogP and logP(o/w) ($\mathrm{|R|}=0.898$).}
%     \label{fig: Figure1}
% \end{figure*}


% \label{note:Note1} 


%TC:endignore
%the command above ignores this section for word count

\end{document}
