## Supplementary material for "CycPeptMP: Enhancing Membrane Permeability Prediction of Cyclic Peptides with Multi-Level Molecular Features and Data Augmentation": latex source: Introduction.tex

\section*{Introduction}
% Cyclic peptides
Cyclic peptides are increasingly valued in pharmaceutical research due to their high binding affinity, target selectivity, and ability to inhibit intracellular protein--protein interactions \cite{Jwad2020, Bhardwaj2022}.
The unique structural features of macrocyclic peptides stem from their restricted conformational flexibility and local secondary structure motifs, which allow for bioactive conformations with remarkable potency and selectivity \cite{Ghosh2023}.
% human peptide hormones: oxytocin
In the past century, cyclic peptide drugs were predominantly sourced from natural products, including antimicrobial agents and human peptide hormones. 
Recent advances in synthesis and screening systems have led to breakthroughs in cyclic peptide drug discovery. 
Novel screening and cyclization strategies have greatly promoted cyclic peptide drug development in the past two decades \cite{Sohrabi2020, Zhang2022}. 
For example, the random nonstandard peptides integrated discovery (RaPID) system designs cyclic peptides from a diverse library including non-natural amino acids, enabling the synthesis and rapid selection of potent binders for therapeutic targets \cite{Yamagishi2011}. 
More than $40$ cyclic peptide drugs are currently in clinical use, and the FDA has approved approximately one macrocyclic peptide drug annually for the past $20$ years \cite{Zorzi2017, Zhang2022}.
% Permeability
Despite their pharmacological potential, cyclic peptides often exhibit poor membrane permeability, severely limiting their biological applications and development of orally available drugs \cite{Vinogradov2019}. 
While the mechanism of membrane permeation by cyclic peptides remains unclear, cyclic peptides with ``closed''-conformation in hydrophobic environments demonstrate enhanced permeability \cite{Whitty2016, Danelius2020}. 
The ``closed'' conformation conceals polar groups through intramolecular hydrogen bonds and lipophilic side chains, contributing to their increased permeation efficiency. 
Drawing inspiration from the structure of Cyclosporin A, a naturally occurring N-methylated macrocyclic peptide with high permeability, the shielding of the exposed hydrogen bond donor (-NH) through N-methylation has been widely applied to enhance membrane permeability \cite{Biron2008, White2011}. 
Various strategies have emerged for improving membrane permeability, such as conformational control \cite{Frost2016, Bhardwaj2022}, amide-to-ester substitution \cite{Hosono2023}, amide-to-thioamide substitution \cite{Ghosh2023}, and side-chain modifications \cite{Taechalertpaisarn2022}. 
However, these strategies do not improve membrane permeability across all cyclic peptides.


% MD & ML based 
In the early stages of drug development, it is important to select candidate compounds with high membrane permeability. 
Because it is costly to randomly measure numerous peptides using biochemical assays, a rapid computational method for predicting membrane permeability is eagerly anticipated. 
Computational approaches for predicting the permeability of cyclic peptides have primarily been based on molecular dynamics (MD) simulation \cite{Witek2019, Ono2019, Sugita2021, Sugita2022}. 
Markov state models, among others, have been used to analyze simulation data and elucidate cyclic peptide behavior \cite{Wang2021}, which is crucial for understanding membrane permeation mechanisms and optimizing structures for enhancing membrane permeability. 
However, the computational cost of MD-based methods is a major limitation.
% Guimaraes2012, 
In contrast to MD-based methods, several physicochemical or machine learning models have been developed, offering more rapid prediction capabilities \cite{Rezai2006, Wang2015, Over2016, Digiesi2021, Poongavanam2021}. 
The descriptors for hydrophobicity, such as the octanol-water partition coefficient (LogP), are generally the most important features for prediction.
% 11, 62, 240?, 62, 20
However, all of these models were established using limited data sets ($10$--$250$ cyclic peptides) and lack a sufficient degree of generalization performance. 
Furthermore, these methods directly apply whole-molecule features used for predicting the membrane permeability of small molecules, ignoring the unique structural characteristics of cyclic peptides, such as sequence information and circularity.


% DL & CycPeptMPDB
Unlike conventional physicochemical and machine learning approaches, deep learning (DL) models offer an architectural design tailored to peptide characteristics \cite{Li2022} and can automatically extract more complex structural features than small-molecule compounds from datasets. 
DL-based small molecule property prediction methods based on graph neural networks (GNNs) and transformers have become a major research area. 
By representing molecule atoms as nodes and bonds as edges, the GNN-based method can effectively capture molecular structural information and integrate the topological structure of molecules with complex atomic features. 
Nonetheless, most existing approaches, such as GCN \cite{Duvenaud2015convolutional, Coley2017convolutional}, GAT \cite{Zhang2021fragat, Choo2023fingerprint}, and MPNN \cite{Withnall2020building}, have intrinsic limitations, such as a poor ability to process global information and high risk of over-smoothing. 
In contrast, following the successful experience in natural language processing, many transformer-based methods have been proposed that treat SMILES as strings \cite{Chithrananda2020chemberta, Zhang2021mg, Irwin2022chemformer}. 
Since these methods lack structural information, several methods have been developed using molecular graph representations as input, which can encode more complex atomic and bond information than strings \cite{Maziarka2020MAT, Chen2022sat, Gao2023transfoxmol, Jiang2023PharmHGT}.
However, cyclic peptide permeability datasets are limited and diverse, and contain discrepancies and errors stemming from the use of various assay systems.
To address these limitations, we constructed CycPeptMPDB \cite{Li2023}: a comprehensive database of cyclic peptide membrane permeability. 
CycPeptMPDB comprises information on a total of $7334$ cyclic peptides, including structures and experimentally measured membrane permeabilities, from $45$ published studies and $2$ patents from pharmaceutical companies, representing the first platform for developing DL-based prediction methods.
Interestingly, over $99.6\%$ of cyclic peptides include non-natural amino acids, which suggests that their creators aimed to enhance permeability through chemical modifications such as N-methylation, or by deliberately incorporating non-natural building blocks in their design.
% In addition to experimental data, the database contains relevant supporting information, such as 3D conformations and hierarchical editing language for macromolecules (HELM) sequence representations, allowing users to analyze data for various applications.


% this study
Cyclic peptides are characterized by complicated conformational dynamics, where even a minor alteration in a single residue can lead to substantial changes in their membrane permeability \cite{Hewitt2015}. 
Therefore, many publications in CycPeptMPDB have focused on measuring changes in membrane permeability while varying only a few residues and maintaining a largely constant sequence. 
In the prediction of small molecule properties \cite{Jiang2023PharmHGT} and peptide--protein binding \cite{Lei2021}, the combined use of multi-scale molecular features can improve prediction accuracy.
This study proposes CycPeptMP: a membrane permeability prediction model for cyclic peptides that effectively integrates multi-level features with state-of-the-art DL techniques. 
We engineered features at the peptide, monomer (substructures, such as residues), and atom levels to concurrently capture both the local sequence variations and global conformational changes in cyclic peptides. 
Additionally, to enhance the training efficiency of our model for complex cyclic peptides, we employed data augmentation methods at the three scales. 
% , provided highly interpretable results, based on various case studies, and effectively identified substructures that significantly influence changes in membrane permeability.
