## Supplementary material for "CycPeptMP: Enhancing Membrane Permeability Prediction of Cyclic Peptides with Multi-Level Molecular Features and Data Augmentation": latex source: Methods.tex

\section*{Materials and Methods}

\subsection*{Experimental dataset}
% CycPeptMPDB PAMPA
We used the structure and membrane permeability ($\mathrm{LogP_{exp}}$) of peptides in CycPeptMPDB. 
CycPeptMPDB contains membrane permeability data based on the parallel artificial membrane permeability (PAMPA), Caco-2, Madin--Darby canine kidney (MDCK), and Ralph Russ canine kidney (RRCK) assays.
% CycPeptMPDB contains $7334$ peptides ($7451$, including duplicated structures from all publications), among which $6941$ are based on the PAMPA, $649$ on the Caco-2 assay, $40$ on the MDCK assay, and $186$ on the RRCK assay. 
% The mechanisms underlying membrane permeation vary considerably between artificial membrane-based assays (i.e., PAMPA) and those using biological membranes. 
We selected PAMPA entries with the largest number of data points. 
If the same peptide was measured in multiple publications, we used the value recorded in the latest publication. 
Consequently, $6889$ peptides were selected, covering an extremely wide range of molecular weights, from $342.44$ to $1777.74$. 
Considering that the lower limit of $\mathrm{LogP_{exp}}$ in CycPeptMPDB was $-10$ ($1\times10^{-10}\mathrm{cm/s}$), but the detection limit in most publications was $-8$ ($1\times10^{-8}\mathrm{cm/s}$), we rounded the permeabilities of $314$ peptides with values lower than $-8$ to $-8$. 
Similarly, the permeability of one peptide with a value higher than $-4$ was rounded to $-4$.


% Split
The validation and test sets were extracted from the overall data for model evaluation. 
First, the Kennard-Stone (KS) algorithm was employed to extract $5\%$ of all data ($344$ peptides) as the test set, which should uniformly cover the multidimensional space \cite{Galvao2005}. 
We generated $2048$-bit Morgan fingerprints (Morgan FP, radius: $2$) and selected the test set such that the Euclidean distance between each data point was maximized by the KS algorithm.
From the remaining data, we randomly extracted $5\%$ three times for the validation set ($344$ peptides), with no overlap between datasets. 
The membrane permeability and molecular weight distributions for each set are shown in Figure~\ref{fig_set-distribution}. 
The average root mean squared error (RMSE) and correlation coefficient (R) from three repeated runs were used as evaluation metrics.


\begin{figure}[tb]
    \centering
    \includegraphics[width=\linewidth, natwidth=14.68cm, natheight=19.47cm]{Fig/set-distribution.png}
    \caption{Experimental data distribution. (A) Membrane permeability. (B) Molecular weight. Valid--1 is the dataset used for the first-time evaluation of the validation set; the corresponding training data sets are Train, Valid--2, and Valid--3.}
    \label{fig_set-distribution}
\end{figure}


\subsection*{Overview of CycPeptMP framework}


\begin{figure*}[tb]
    \centering
    \includegraphics[width=\linewidth, natwidth=40.22cm, natheight=19.93cm]{Fig/framework.png}
    \caption{Overall framework of CycPeptMP model. The model incorporates the Transformer-based atom, convolutional neural network (CNN)-based monomer, and MLP-based peptide sub-models. The three-level expression vectors extracted by the three sub-models are concatenated and passed through a shared layer to derive the final permeability prediction value.}
    \label{fig_framework}
\end{figure*}


Figure~\ref{fig_framework} shows the overall architecture of the CycPeptMP model. 
We designed three-level peptide representations and used each for three different sub-models to extract the atom-, monomer-, and peptide-level molecular representations. 
First, the input peptide was divided into monomers, and respective 3D conformations were generated from the peptide and monomers. 
Next, atom- and peptide-level features were extracted from the peptide conformation and used as input for the atom and peptide models, respectively. 
Monomer-level features extracted from the monomer conformation were used as input for the monomer model. 
Finally, the three-level latent feature vectors extracted using the three sub-models were combined to derive the membrane permeability prediction value.


\subsection*{Division of the monomer}
We designed monomer-level features to accurately capture the subtle structural variations of cyclic peptides. 
The definition of a monomer corresponds to that provided by CycPeptMPDB. 
Although the peptide and ester bonds on the side chain were cleaved in CycPeptMPDB, bonds existing anywhere other than the macrocycle were not subjected to division to completely express the properties of the local structure.
Merely hydrolyzing the peptide bond could generate a new hydrogen bond donor, which could misrepresent the original physicochemical properties of the substructure. 
Hence, when generating the conformation and calculating the monomer descriptor, the cleaved amide group or O atom of the amide-to-ester substitution was methylated (addition of CH3), and the carboxyl group was converted to an aldehyde group (addition of H).


\subsection*{Peptide and monomer descriptors}

To design peptide- and monomer-level features, peptides and monomers were represented by $16$ descriptors, including LogP and polar surface area (Table~\ref{tab_descriptors}, the correlation matrix of the selected descriptors is shown in \SupFigDesccorr). 
First, the 3D conformations of peptides and monomers were generated using RDKit. 
The initial structures were generated using the ETKDG method and then optimized using the UFF force field. 
Then, the 2D and 3D descriptors were calculated; we used a single-conformation 3D structure in the calculation.
The $16$ descriptors were selected as follows: 
First, a total of $1857$ descriptors ($1689$ 2D and $168$ 3D descriptors) were calculated for both cyclic peptides (peptide descriptors) and each monomer (monomer descriptors) using MOE software (version 2019.01) \cite{MOE}, the Mordred package (version 1.2) \cite{Moriwaki2018}, and RDKit package (version 2022.09.5) \cite{RDKit}. 
For 2D and 3D peptide descriptors, we removed all descriptors with constant values among cyclic peptides.
For descriptor pairs with an absolute correlation coefficient of $0.9$ or more, we excluded the one with the lower correlation with permeability. 
Consequently, $407$ ($335$ 2D and $72$ 3D descriptors) peptide descriptors were selected.
We constructed two random forest (RF) models for predicting permeability based on the 2D or 3D peptide descriptors. 
Seven 2D and nine 3D peptide descriptors were selected based on the feature importance determined by the RF models (\SupFigDescpre). 
Finally, the same $16$ monomer descriptors were selected, and both peptide and monomer descriptors were standardized based on the Z-score.


\begin{table*}[tb]
\caption{Selected descriptors, arranged in order of importance.}
\label{tab_descriptors}
\centering
\scalebox{0.75}{
\begin{tabular}{llll}
\toprule
Type & Name           & Calculated by & Description                                                                                                      \\
\midrule
2D   & VSA\_EState9   & Mordred       & Van der Waals surface area using EState indices and surface area contribution                                    \\
     & density        & MOE           & Molecular mass density (molecular weight divided by van der Waals volume using a connection table approximation) \\
     & MolLogP        & RDKit         & Wildman--Crippen LogP value                                                                                      \\
     & fr\_Al\_OH     & RDKit         & Number of aliphatic hydroxyl groups                                                                              \\
     & logP(o/w)      & MOE           & Log of the octanol/water partition coefficient                                                                   \\
     & lip\_violation & MOE           & Number of violations of Lipinski's Rule of Five                                                                  \\
     & h\_logD        & MOE           & Log of the octanol/water distribution coefficient at pH 7                                                        \\
3D   & dens           & MOE           & Molecular mass density (molecular weight divided by 3D van der Waals volume)                                     \\
     & FNSA4          & Mordred       & Fractional charged partial negative surface area (version 4)                                                     \\
     & RNCS           & Mordred       & Relative negative charge surface area                                                                            \\
     & FASA-          & MOE           & Fractional water accessible surface area of all atoms with negative partial charge                               \\
     & FCASA+         & MOE           & Fractional positive charge weighted surface area                                                                 \\
     & FASA\_P        & MOE           & Fractional water accessible surface area of all polar atoms                                                      \\
     & FNSA2          & Mordred       & Fractional charged partial negative surface area (version 2)                                                     \\
     & FNSA5          & Mordred       & Fractional charged partial negative surface area (version 5)                                                     \\
     & vsurf\_Wp2     & MOE           & VolSurf polar volume                                                                                             \\
\bottomrule
\end{tabular}
}
\end{table*}


\begin{figure*}[tb]
    \centering
    \includegraphics[width=\linewidth, natwidth=30.2cm, natheight=10.69cm]{Fig/model.png}
    \caption{(A) Architecture of atom model. Node features and three types of bond matrices were used as input for the Transformer-based model. (B) Architecture of the monomer model. Monomer descriptors were used as input for the CNN-based model.}
    \label{fig_model}
\end{figure*}


\subsection*{Atom model}


\begin{table*}[tb]
\caption{Input features of atom model.}
\label{tab_atom-features}
\centering
\scalebox{0.9}{
\begin{tabular}{llrl}
\toprule
Type  & Feature                    & Size & Description                                                                          \\
\midrule
Atom  & Atom type                  & 8    & Type of atom by atomic number (one-hot)                                              \\
      & Degree of atom             & 5    & Number of bonds (one-hot)                                                            \\
      & Number of hydrogen         & 5    & Number of bonded hydrogen atoms (one-hot)                                            \\
      & Formal charge              & 4    & Electrical charge (one-hot)                                                           \\
      & Hybridization              & 3    & Type of atom hybridization (one-hot)                                                  \\
      & Chirality                  & 3    & Type of atom chirality (one-hot)                                                      \\
      & Is in a ring               & 1    & Whether included in a ring structure                                                  \\
      & Is aromatic                & 1    & Whether aromatic                                                                      \\
Bond  & Bond type ($Bond$)         & 1    & Single(1.0), Double(2.0), Triple(3.0), Aromatic(1.5), Conjugated(1.4) and No-bond (0) \\
      & Graph distance ($Graph$)   & 1    & Distance calculated from graph representation                                         \\
      & 3D distance ($Conf$)       & 1    & Euclidean distance (\AA) calculated from 3D conformation                              \\
\bottomrule
\end{tabular}
}
\end{table*}


We constructed a transformer-based atom model to capture the overall graph structure and 3D conformation of the peptide (Figure~\ref{fig_model}~(A)). 
We used node features ($Node$) and three types of bond weight matrices as inputs for the atom model ($Bond, Graph, Conf$).
As shown in Table~\ref{tab_atom-features}, heavy atoms were considered nodes and node features were represented as $Node$, bonded interaction weights were represented as $Bond \in \mathbb{R}^{N_{atoms} \times N_{atoms}}$, graphic pairwise distances were represented as $Graph \in \mathbb{R}^{N_{atoms} \times N_{atoms}}$, and 3D pairwise distances were represented as $Conf \in \mathbb{R}^{N_{atoms} \times N_{atoms}}$.


$Bond$ recorded the molecule bond information and controlled message propagation between neighboring nodes by assigning weights to each bond type. 
According to the chemical bonding principle, bonds with more electron participation, such as unsaturated bonds, were assigned higher weights to enhance the exchange of information between atoms \cite{Gao2023transfoxmol}. 
To capture the local relationship between each node, embedded $Bond$ was used for positional encoding and added to embedded $Node$ to serve as the input for the encoder block as follows:
\begin{equation}
h = \frac{W_{node} Node + W_{bond} Bond}{\sqrt{d_{model}}}
\label{equ_input}
\end{equation}
Where $d_{model}$ is the attention dimension of the atom model, $W_{node} \in \mathbb{R}^{N_{node-features} \times d_{model}}$ and $W_{bond} \in \mathbb{R}^{N_{atoms} \times d_{model}}$ are trainable parameters, and $h \in \mathbb{R}^{N_{atoms} \times d_{model}}$ is the updated input for the encoder block.


Two types of distance matrices, $Graph$ and $Conf$, the shortest pairwise graph distance and 3D Euclidean distance of each atom, were used to capture the overall structure and 3D conformation of the peptide. 
The distance maps were processed through an attenuation function to weaken distant interactions as follows:
\begin{subequations}
\label{equ_distance}
\begin{align}
Strength^{graph}_{i, j} &= 
\left\{
    \begin{array}{cc}
    1 & (i = j) \\
    \frac{1}{Graph_{i, j}} & (i \ne j)
    \end{array}
\right. \\
Strength^{conf}_{i, j} &= 
\left\{
    \begin{array}{cc}
    1 & (i = j) \\
    \frac{1}{Conf_{i, j}} & (i \ne j)
    \end{array}
\right.
\end{align}
\end{subequations}
Here, we designed a structure-enhanced transformer encoder to learn the structural and 3D conformational information of peptides using focused attention. 
The encoder consisted of two parts, one using $Strength^{graph}$ and another using $Strength^{conf}$. 
This approach attenuates attention between less relevant pairs based on the distance, as shown in Equation~(\ref{equ_attention_graph}) and Equation~(\ref{equ_attention_conf}), providing a simplified approach to modeling complex molecular structures.
\begin{subequations}
\label{equ_attention_graph}
\begin{align}
\mathrm{Atten}(h^{graph^{l-1}}) & = \mathrm{softmax}\left(\frac{Q K^{T}}{\sqrt{d_{model}}}\right)V \\
h^{graph^{l}}     & = \mathrm{Atten}(h^{graph^{l-1}}) \odot Strength^{graph} \\
h^{graph^{l}}     & = \mathrm{LayerNorm}(h^{graph^{l-1}} + h^{graph^{l}}) \\
h^{graph^{l}}     & = \mathrm{LayerNorm}(h^{graph^{l}} + \mathrm{FFN}(h^{graph^{l}}))
\end{align}
\end{subequations}
\begin{subequations}
\label{equ_attention_conf}
\begin{align}
\mathrm{Atten}(h^{conf^{l-1}}) & = \mathrm{softmax}\left(\frac{Q K^{T}}{\sqrt{d_{model}}}\right)V \\
h^{conf^{l}}     & = \mathrm{Atten}(h^{conf^{l-1}}) \odot Strength^{conf} \\
h^{conf^{l}}     & = \mathrm{LayerNorm}(h^{conf^{l-1}} + h^{conf^{l}}) \\
h^{conf^{l}}     & = \mathrm{LayerNorm}(h^{conf^{l}} + \mathrm{FFN}(h^{conf^{l}}))
\end{align}
\end{subequations}
where $h^{graph^{l-1}}$ and $h^{graph^{l}}$ are the updated latent features of the $Graph$ block ($l-1$)-th and $l$-th layers, respectively, and $h^{conf^{l-1}}$ and $h^{conf^{l}}$ are the updated latent features of the $Conf$ block ($l-1$)-th and $l$-th layers, respectively.


Finally, the outputs $h^{graph^{out}}$ and $h^{conf^{out}}$ of the two blocks were weighted using the hyperparameter $\lambda_{g}$ and the concatenated feature vector was used to derive the final output $out^{atom}$ of the atom model as follows:
\begin{equation}
out^{atom} = \mathrm{Linear}(\mathrm{Concat}(\lambda_{g} * h^{graph^{out}}, (1 - \lambda_{g}) * h^{conf^{out}}))
\label{equ_outatom}
\end{equation}


\subsection*{Monomer model}
To capture the partial structural information of peptides at the sequence level, we constructed a monomer model based on the $16$ types of monomer descriptors (Figure~\ref{fig_model}~(B)). 
The CNN was used to learn partial structural features and sequence information.
% We thought the CNN model would be suitable for learning the partial structural features and sequence information. However, the general 1D-CNN layer cannot express the circularity of cyclic peptides, and the input is treated as a linear peptide. Therefore, we incorporated the CyclicConv layer \cite{Li2022}, which supplemented adjacent monomers at both ends of an input peptide sequence before performing convolution.
For the convolution layer, we used the general 1D-CNN layer or a CyclicConv layer \cite{Li2022} that considers peptide circularity. 
The use of the CyclicConv or 1D-CNN layer was determined by hyperparameter tuning. 
Finally, the monomer model derived the latent feature $out^{monomer}$.


\subsection*{Peptide model}
The $16$ peptide descriptors all expressed the physicochemical properties of the molecules. 
We also incorporated $2048$-bit Morgan FP ($1024$-bit, radius: $2$; $1024$-bit, radius: $3$) to express the substructural information for the peptide model. 
The descriptor and Morgan FP were each trained with different multilayer perceptrons (MLPs), and the latent feature vectors $h^{desc^{out}}$ and $h^{fp^{out}}$ were concatenated and used to derive the final output $out^{peptide}$ of the peptide model as follows:
\begin{equation}
out^{peptide} = \mathrm{Linear}(\mathrm{Concat}(h^{desc^{out}}, h^{fp^{out}}))
\label{equ_outpep}
\end{equation}


\subsection*{Fusion model}
As shown in Figure~\ref{fig_framework}, the output latent feature vectors $out^{atom}$, $out^{monomer}$, and $out^{peptide}$ of the three sub-models were concatenated to generate the final molecular feature vector, which was passed through a shared layer for the final permeability prediction $out^{fusion}$. 
As the model becomes more complex, problems such as gradient disappearance may occur and input information may not be transmitted. 
Auxiliary loss is a learning technique in which additional losses are incurred to optimize the NN learning process. 
Directly propagating errors to the middle network layer can prevent gradient disappearance and improve embedding and learning efficiency \cite{Zhou2019deep}.
Hence, in addition to the main loss $L$ calculated from the output of the fusion model, we designed the three sub-model losses $L_{atom}$, $L_{monomer}$, and $L_{peptide}$ derived from the output of each sub-model, and layer losses $L_{layer_{a}}$, $L_{layer_{m}}$, and $L_{layer_{p}}$ derived from the averaged outputs of the layers in each block (Transformer, CNN, and MLP) of the three sub-models. 
The loss function during training is presented in Equation~(\ref{equ_aux}):
\begin{equation}
\begin{split}
Loss = L &+ \gamma_{sub} * (L_{atom} + L_{monomer} + L_{peptide}) \\
         &+ \gamma_{layer} * (L_{layer_{a}} + L_{layer_{m}} + L_{layer_{p}})
\label{equ_aux}
\end{split}
\end{equation}
where the weight parameter $\gamma_{sub}$ was set to $0.10$ and $\gamma_{layer}$ was set to $0.05$. 
Only the output value $out^{fusion}$ of the fusion model was used during inference.
The hyperparameters of CycPeptMP were determined by $150$ searches using Optuna software \cite{Akiba2019optuna}; the search range and results are shown in \SupTabOptuna.


 

\subsection*{Data augmentation}


\begin{figure}[tb]
    \centering
    \includegraphics[width=\linewidth, natwidth=25.54cm, natheight=16.65cm]{Fig/augmentation.png}
    \caption{Sequence arrangement in the monomer model. The aligned monomer descriptors are translated and rotated based on the sequence information.}
    \label{fig_augmentation}
\end{figure}


Although the amount of available biological data has increased in recent years, experimental data remains limited compared to data for natural language processing and computer vision. 
For example, the Tox21 dataset, which deals with the toxicity classification of small molecules, only has about $8000$ data points. 
This limitation in biological data has motivated the increased use of self-supervised learning approaches utilizing contrastive learning \cite{Wang2022molecular} and pre-training \cite{Chithrananda2020chemberta, Irwin2022chemformer}. 
However, given a more limited availability compared to small molecules, these techniques remain challenging for cyclic peptides.
To improve the learning efficiency of the model, we used three augmentation methods to generate $60$ replicas based on the properties of SMILES, nature of cyclic peptide sequences, and the complexity of structural changes in cyclic peptides. 
First, the SMILES enumeration technique \cite{Bjerrum2017smiles} was used to permute the atom order and generate input for the atom model with a different ordering. 
Next, considering the circularity of the cyclic peptide, the input of the monomer model was rearranged using sequence arrangement---the aligned monomer descriptors were translated and rotated as shown in Figure~\ref{fig_augmentation}. 
Finally, $60$ conformations per peptide/monomer were generated using RDKit to incorporate more diverse 3D information into the model.
Cyclic peptide conformations were used to calculate the $Conf$ matrix for the atom model and peptide descriptors for the peptide model, and monomer conformations were used to calculate the monomer descriptors for the monomer model. 
During training, each replica was given the same label and treated as independent data. 
During inference, the average of the $60$ replicas was used as the final predicted value.


\subsection*{Baseline methods}
We validated the performance of CycPeptMP based on comparisons with seven baseline methods.
\begin{itemize}
\item 
Traditional baselines: 
To represent traditional cyclic peptide membrane permeability prediction methods, we constructed a RF model with $2048$-bit Morgan FP, support vector machine (SVM) model with seven 2D peptide descriptors, and SVM model with $16$ 2D and 3D peptide descriptors. 
The hyperparameters of the RF and SVM models were determined by a grid search (\SupTabGrid).
\item
Transformer-based methods:
MAT \cite{Maziarka2020MAT} and SAT \cite{Chen2022sat} were compared as state-of-the-art transformer-based methods for predicting small-molecule properties. 
MAT is a graph-based transformer that encodes the graph topology by adding weighted adjacency and 3D distance matrices to the self-attention. 
SAT is a structure-aware transformer that incorporates structural information into the original self-attention by extracting a subgraph representation rooted at each node before computing the attention.
\item
Multi-level feature methods:
PharmHGT \cite{Jiang2023PharmHGT} designs features on the atom- and fragment-level and constructs a heterogeneous graph considering the correspondence between atoms and fragments for a transformer-based model. 
FinGAT \cite{Choo2023fingerprint} uses a GAT model to extract atom-level information and combines it with Morgan FP to capture the molecular structure from multiple perspectives.
\end{itemize}
The same validation set, test set, and random seeds of the DL-based model were used for all models to ensure a fair comparison. 
