## Supplementary material for "CycPeptMP: Enhancing Membrane Permeability Prediction of Cyclic Peptides with Multi-Level Molecular Features and Data Augmentation": latex source: reference.bib

\bibitem{bahdanau2014neural}
Dzmitry Bahdanau, Kyunghyun Cho, and Yoshua Bengio
\newblock Neural machine translation by jointly learning to align and
  translate
\newblock {\em arXiv preprint arXiv:14090473}, 2014

@article{Bhardwaj2022,
    author = {Bhardwaj, Gaurav and O'Connor, Jacob and Rettie, Stephen and Huang, Yen-Hua and Ramelot, Theresa A and Mulligan, Vikram Khipple and Alpkilic, Gizem Gokce and Palmer, Jonathan and Bera, Asim K and Bick, Matthew J and {Di Piazza}, Maddalena and Li, Xinting and Hosseinzadeh, Parisa and Craven, Timothy W and Tejero, Roberto and Lauko, Anna and Choi, Ryan and Glynn, Calina and Dong, Linlin and Griffin, Robert and van Voorhis, Wesley C and Rodriguez, Jose and Stewart, Lance and Montelione, Gaetano T and Craik, David and Baker, David},
    journal = {Cell},
    month = {aug},
    pages = {1--13},
    title = {{Accurate de novo design of membrane-traversing macrocycles}},
    year = {2022}
}

@article{Jwad2020,
    title={{Strategies for fine-tuning the conformations of cyclic peptides}},
    author={Jwad, Rasha and Weissberger, Daniel and Hunter, Luke},
    journal={Chem Rev},
    volume={120},
    number={17},
    pages={9743--9789},
    year={2020},
}

@article{Ghosh2023,
    author = {Ghosh, Pritha and Raj, Nishant and Verma, Hitesh and Patel, Monika and Chakraborti, Sohini and Khatri, Bhavesh and Doreswamy, Chandrashekar M and Anandakumar, S R and Seekallu, Srinivas and Dinesh, M B and Jadhav, Gajanan and Yadav, Prem Narayan and Chatterjee, Jayanta},
    journal={Nat Commun},
    number = {1},
    title = {{An amide to thioamide substitution improves the permeability and bioavailability of macrocyclic peptides}},
    volume = {14},
    year = {2023}
}


@article{Zorzi2017,
    author = {Zorzi, Alessandro and Deyle, Kaycie and Heinis, Christian},
    title = {{Cyclic peptide therapeutics: past, present and future}},
    journal = {Curr Opin Chem Biol},
    year = {2017},
    volume = {38},
    pages = {24--29},
}

@article{Sohrabi2020,
    author = {Sohrabi, Catrin and Foster, Andrew and Tavassoli, Ali},
    journal = {Nat Rev Chem},
    number = {2},
    pages = {90--101},
    title = {{Methods for generating and screening libraries of genetically encoded cyclic peptides in drug discovery}},
    volume = {4},
    year = {2020},
}

@article{Yamagishi2011,
    title={{Natural product-like macrocyclic N-methyl-peptide inhibitors against a ubiquitin ligase uncovered from a ribosome-expressed de novo library}},
    author={Yamagishi, Yusuke and Shoji, Ikuo and Miyagawa, Shoji and Kawakami, Takashi and Katoh, Takayuki and Goto, Yuki and Suga, Hiroaki},
    journal={Chem Biol},
    volume={18},
    number={12},
    pages={1562--1570},
    year={2011},
}

@article{Vinogradov2019,
    author = {Vinogradov, Alexander A and Yin, Yizhen and Suga, Hiroaki},
    title = {{Macrocyclic Peptides as Drug Candidates: Recent Progress and Remaining Challenges}},
    journal = {J Am Chem Soc},
    year = {2019},
    volume = {141},
    number = {10},
    pages = {4167--4181},
}

@article{Zhang2022,
    title={{Cyclic peptide drugs approved in the last two decades (2001--2021)}},
    author={Zhang, Huiya and Chen, Shiyu},
    journal={RSC Chem Biol},
    volume={3},
    number={1},
    pages={18--31},
    year={2022},
}

@article{Whitty2016,
    title={{Quantifying the chameleonic properties of macrocycles and other high-molecular-weight drugs}},
    author = {Whitty, Adrian and Zhong, Mengqi and Viarengo, Lauren and Beglov, Dmitri and Hall, David R and Vajda, Sandor},
    journal={Drug Discov Today},
    volume={21},
    number={5},
    pages={712--717},
    year={2016},
}

@article{Danelius2020,
    title={{Solution conformations explain the chameleonic behaviour of macrocyclic drugs}},
    author = {Danelius, Emma and Poongavanam, Vasanthanathan and Peintner, Stefan and Wieske, Lianne HE and Erd{\'{e}}lyi, M{\'{a}}t{\'{e}} and Kihlberg, Jan},
    journal={Chem Eur J},
    volume={26},
    number={23},
    pages={5231--5244},
    year={2020},
}

@article{Biron2008,
    title={{Improving oral bioavailability of peptides by multiple N-methylation: somatostatin analogues}},
    author = {Biron, Eric and Chatterjee, Jayanta and Ovadia, Oded and Langenegger, Daniel and Brueggen, Joseph and Hoyer, Daniel and Schmid, Herbert A and Jelinek, Raz and Gilon, Chaim and Hoffman, Amnon and Kessler, Horst},
    journal={Angew Chem Int Ed},
    volume={47},
    number={14},
    pages={2595--2599},
    year={2008},
}


@article{Frost2016,
    title={{Oxadiazole grafts in peptide macrocycles}},
    author={Frost, John R and Scully, Conor C G and Yudin, Andrei K},
    journal={Nat Chem},
    volume={8},
    number={12},
    pages={1105--1111},
    year={2016},
}

@article{White2011,
    title={{On-resin N-methylation of cyclic peptides for discovery of orally bioavailable scaffolds}},
    author = {White, Tina R and Renzelman, Chad M and Rand, Arthur C and Rezai, Taha and McEwen, Cayla M and Gelev, Vladimir M and Turner, Rushia A and Linington, Roger G and Leung, Siegfried SF and Kalgutkar, Amit S and Bauman, Jonathan N and Zhang, Yizhong and Liras, Spiros and Price, David A and Mathiowetz, Alan M and Jacobson, Matthew P and Lokey, R Scott},
    journal={Nat Chem Biol},
    volume={7},
    number={11},
    pages={810--817},
    year={2011},
}


@article{Hosono2023,
    title={{Amide-to-ester substitution as a stable alternative to N-methylation for increasing membrane permeability in cyclic peptides}},
    author = {Hosono, Yuki and Uchida, Satoshi and Shinkai, Moe and Townsend, Chad E and Kelly, Colin N and Naylor, Matthew R and Lee, Hsiau-wei and Kanamitsu, Kayoko and Ishii, Mayumi and Ueki, Ryosuke and Ueda, Takumi and Takeuchi, Koh and Sugita, Masatake and Akiyama, Yutaka and Lokey, Scott R and Morimoto, Jumpei and Sando, Shinsuke},
    journal={Nat Commun},
    volume={14},
    number={1},
    pages={1416},
    year={2023},
}

@article{Taechalertpaisarn2022,
    title={{A new amino acid for improving permeability and solubility in macrocyclic peptides through side chain-to-backbone hydrogen bonding}},
    author={Taechalertpaisarn, Jaru and Ono, Satoshi and Okada, Okimasa and Johnstone, Timothy C and Lokey, R Scott},
    journal={J Med Chem},
    volume={65},
    number={6},
    pages={5072--5084},
    year={2022},
}

@article{Witek2019,
    title={{Rationalization of the membrane permeability differences in a series of analogue cyclic decapeptides}},
    author = {Witek, Jagna and Wang, Shuzhe and Schroeder, Benjamin and Lingwood, Robin and Dounas, Andreas and Roth, Hans J{\"{o}}rg and Fouch{\'{e}}, Marianne and Blatter, Markus and Lemke, Oliver and Keller, Bettina and Riniker, Sereina},
    journal={J Chem Inf Model},
    volume={59},
    number={1},
    pages={294--308},
    year={2019},
}

@article{Ono2019,
    title={{Conformation and permeability: cyclic hexapeptide diastereomers}},
    author={Ono, Satoshi and Naylor, Matthew R and Townsend, Chad E and Okumura, Chieko and Okada, Okimasa and Lokey, R Scott},
    journal={J Chem Inf Model},
    volume={59},
    number={6},
    pages={2952--2963},
    year={2019},
    publisher={ACS Publications}
}


@article{Sugita2021,
    title={{Large-scale membrane permeability prediction of cyclic peptides crossing a lipid bilayer based on enhanced sampling molecular dynamics simulations}},
    author={Sugita, Masatake and Sugiyama, Satoshi and Fujie, Takuya and Yoshikawa, Yasushi and Yanagisawa, Keisuke and Ohue, Masahito and Akiyama, Yutaka},
    journal={J Chem Inf Model},
    volume={61},
    number={7},
    pages={3681--3695},
    year={2021},
}

@article{Sugita2022,
    title={{Lipid composition is critical for accurate membrane permeability prediction of cyclic peptides by molecular dynamics simulations}},
    author={Sugita, Masatake and Fujie, Takuya and Yanagisawa, Keisuke and Ohue, Masahito and Akiyama, Yutaka},
    journal={J Chem Inf Model},
    volume={62},
    number={18},
    pages={4549--4560},
    year={2022},
}

@article{Wang2021,
  title={{Effect of flexibility, lipophilicity, and the location of polar residues on the passive membrane permeability of a series of cyclic decapeptides}},
  author={Wang, Shuzhe and Konig, Gerhard and Roth, Hans-Jorg and Fouch{\'e}, Marianne and Rodde, Stephane and Riniker, Sereina},
  journal={J Med Chem},
  volume={64},
  number={17},
  pages={12761--12773},
  year={2021},
}

@article{Rezai2006,
    title={{Conformational flexibility, internal hydrogen bonding, and passive membrane permeability: successful in silico prediction of the relative permeabilities of cyclic peptides}},
    author={Rezai, Taha and Bock, Jonathan E and Zhou, Mai V and Kalyanaraman, Chakrapani and Lokey, R Scott and Jacobson, Matthew P},
    journal = {J Am Chem Soc},
    volume={128},
    number={43},
    pages={14073--14080},
    year={2006},
}

@article{Guimaraes2012,
    title={{Use of 3D properties to characterize beyond rule-of-5 property space for passive permeation}},
    author={Guimar\~{a}es, Cristiano RW and Mathiowetz, Alan M and Shalaeva, Marina and Goetz, Gilles and Liras, Spiros},
    journal={J Chem Inf Model},
    volume={52},
    number={4},
    pages={882--890},
    year={2012},
}

@article{Wang2015,
    title={{Exploring experimental and computational markers of cyclic peptides: Charting islands of permeability}},
    author={Wang, Conan K and Northfield, Susan E and Swedberg, Joakim E and Colless, Barbara and Chaousis, Stephanie and Price, David A and Liras, Spiros and Craik, David J},
    journal={Eur J Med Chem},
    volume={97},
    pages={202--213},
    year={2015},
}


@article{Over2016,
    title={{Structural and conformational determinants of macrocycle cell permeability}},
    author = {Over, Bj{\"{o}}rn and Matsson, P{\"{a}}r and Tyrchan, Christian and Artursson, Per and Doak, Bradley C and Foley, Michael A and Hilgendorf, Constanze and Johnston, Stephen E and Lee, Maurice D and Lewis, Richard J and McCarren, Patrick and Muncipinto, Giovanni and Norinder, Ulf and Perry, Matthew WD and Duvall, Jeremy R and Kihlberg, Jan},
    journal={Nat Chem Biol},
    volume={12},
    number={12},
    pages={1065--1074},
    year={2016},
}

@article{Digiesi2021,
    title={{Permeability prediction in the beyond-Rule-of 5 chemical space: Focus on cyclic hexapeptides}},
    author={Digiesi, Vito and de la Oliva Roque, V{\'\i}ctor and Vallaro, Maura and Caron, Giulia and Ermondi, Giuseppe},
    journal={Eur J Pharm Biopharm},
    volume={165},
    pages={259--270},
    year={2021},
}

@article{Poongavanam2021,
    title={{Predicting the permeability of macrocycles from conformational sampling--limitations of molecular flexibility}},
    author={Poongavanam, Vasanthanathan and Atilaw, Yoseph and Ye, Sofie and Wieske, Lianne HE and Erdelyi, Mate and Ermondi, Giuseppe and Caron, Giulia and Kihlberg, Jan},
    journal={J Pharm Sci},
    volume={110},
    number={1},
    pages={301--313},
    year={2021},
}

@article{Li2022,
    title={{Plasma protein binding prediction focusing on residue-level features and circularity of cyclic peptides by deep learning}},
    author={Li, Jianan and Yanagisawa, Keisuke and Yoshikawa, Yasushi and Ohue, Masahito and Akiyama, Yutaka},
    journal={Bioinformatics},
    volume={38},
    number={4},
    pages={1110--1117},
    year={2022},
}

@article{Li2023,
    title={{CycPeptMPDB: A Comprehensive Database of Membrane Permeability of Cyclic Peptides}},
    author={Li, Jianan and Yanagisawa, Keisuke and Sugita, Masatake and Fujie, Takuya and Ohue, Masahito and Akiyama, Yutaka},
    journal={J Chem Inf Model},
    volume={63},
    number={7},
    pages={2240--2250},
    year={2023},
}

@article{Hewitt2015,
    author = {Hewitt, William M and Leung, Siegfried SF and Pye, Cameron R and Ponkey, Alexandra R and Bednarek, Maria and Jacobson, Matthew P and Lokey, R Scott},
    journal = {J Am Chem Soc},
    number = {2},
    pages = {715--721},
    title = {{Cell-permeable cyclic peptides from synthetic libraries inspired by natural products}},
    volume = {137},
    year = {2015}
}


@article{Jiang2023PharmHGT,
  title={{Pharmacophoric-constrained heterogeneous graph transformer model for molecular property prediction}},
  author={Jiang, Yinghui and Jin, Shuting and Jin, Xurui and Xiao, Xianglu and Wu, Wenfan and Liu, Xiangrong and Zhang, Qiang and Zeng, Xiangxiang and Yang, Guang and Niu, Zhangming},
  journal={Commun Chem},
  volume={6},
  number={1},
  pages={60},
  year={2023},
}

@article{Lei2021,
  title={{A deep-learning framework for multi-level peptide--protein interaction prediction}},
  author={Lei, Yipin and Li, Shuya and Liu, Ziyi and Wan, Fangping and Tian, Tingzhong and Li, Shao and Zhao, Dan and Zeng, Jianyang},
  journal={Nat Commun},
  volume={12},
  number={1},
  pages={5465},
  year={2021},
}

%%%%%%%%%%%%%%%%%%%%%%%%%%%%%%%%%%%%%%%%%%%%%%%%%%%%%%%%%%%%%%%%%%%%%%%%%

@article{Galvao2005,
    title={{A method for calibration and validation subset partitioning}},
    author={Galvao, Roberto Kawakami Harrop and Araujo, M{\'a}rio C{\'e}sar Ugulino and Jos{\'e}, Gledson Em{\'\i}dio and Pontes, Marcio Jos{\'e} Coelho and Silva, Edvan Cirino and Saldanha, Teresa Cristina Bezerra},
    journal={Talanta},
    volume={67},
    number={4},
    pages={736--740},
    year={2005},
}

@manual{MOE,
  author = {Chemical Computing Group Inc.},
  year = {2019},
  title = {{Molecular Operating Environment (MOE), 2019.01}},
  address = {Montreal, QC, Canada},
}

@article{Moriwaki2018,
  title={{Mordred: a molecular descriptor calculator}},
  author={Moriwaki, Hirotomo and Tian, Yu-Shi and Kawashita, Norihito and Takagi, Tatsuya},
  journal={J Cheminform},
  volume={10},
  number={1},
  pages={1--14},
  year={2018},
}


@manual{RDKit,
  author = {Greg Landrum},
  year = {2022},
  title = {{RDKit: Open-source cheminformatics. https://www.rdkit.org}},
}

%%%%%%%%%%%%%%%%%%%%%%%%%%%%%%%%%%%%%%%%%%%%%%%%%%%%%%%%%%%%%%%%%%%%%%%%%

@article{Duvenaud2015convolutional,
  title={{Convolutional networks on graphs for learning molecular fingerprints}},
  author={Duvenaud, David K and Maclaurin, Dougal and Iparraguirre, Jorge and Bombarell, Rafael and Hirzel, Timothy and Aspuru-Guzik, Al{\'a}n and Adams, Ryan P},
  journal={Adv Neural Inf Process Syst},
  volume={28},
  year={2015}
}


@article{Coley2017convolutional,
  title={{Convolutional embedding of attributed molecular graphs for physical property prediction}},
  author={Coley, Connor W and Barzilay, Regina and Green, William H and Jaakkola, Tommi S and Jensen, Klavs F},
  journal={J Chem Inf Model},
  volume={57},
  number={8},
  pages={1757--1772},
  year={2017},
}

@article{Zhang2021fragat,
  title={{FraGAT: a fragment-oriented multi-scale graph attention model for molecular property prediction}},
  author={Zhang, Ziqiao and Guan, Jihong and Zhou, Shuigeng},
  journal={Bioinformatics},
  volume={37},
  number={18},
  pages={2981--2987},
  year={2021},
  publisher={Oxford University Press}
}

@article{Choo2023fingerprint,
  title={{Fingerprint-Enhanced Graph Attention Network (FinGAT) Model for Antibiotic Discovery}},
  author={Choo, Hou Yee and Wee, JunJie and Shen, Cong and Xia, Kelin},
  journal={J Chem Inf Model},
  year={2023},
  volume = {63},
  number = {10},
  pages = {2928--2935},
  publisher={ACS Publications}
}


@article{Withnall2020building,
  title={{Building attention and edge message passing neural networks for bioactivity and physical--chemical property prediction}},
  author={Withnall, Michael and Lindel{\"o}f, Edvard and Engkvist, Ola and Chen, Hongming},
  journal={J Cheminform},
  volume={12},
  number={1},
  pages={1--18},
  year={2020},
  publisher={BioMed Central}
}


@article{Chithrananda2020chemberta,
  title={{ChemBERTa: large-scale self-supervised pretraining for molecular property prediction}},
  author={Chithrananda, Seyone and Grand, Gabriel and Ramsundar, Bharath},
  journal={arXiv preprint arXiv:2010.09885},
  year={2020}
}

@article{Zhang2021mg,
  title={{MG-BERT: leveraging unsupervised atomic representation learning for molecular property prediction}},
  author={Zhang, Xiao-Chen and Wu, Cheng-Kun and Yang, Zhi-Jiang and Wu, Zhen-Xing and Yi, Jia-Cai and Hsieh, Chang-Yu and Hou, Ting-Jun and Cao, Dong-Sheng},
  journal={Brief Bioinform},
  volume={22},
  number={6},
  pages={bbab152},
  year={2021},
}

@article{Irwin2022chemformer,
  title={{Chemformer: a pre-trained transformer for computational chemistry}},
  author={Irwin, Ross and Dimitriadis, Spyridon and He, Jiazhen and Bjerrum, Esben Jannik},
  journal={Mach learn: sci technol},
  volume={3},
  number={1},
  pages={015022},
  year={2022},
}


@article{Maziarka2020MAT,
  title={{Molecule attention transformer}},
  author={Maziarka, {\L}ukasz and Danel, Tomasz and Mucha, S{\l}awomir and Rataj, Krzysztof and Tabor, Jacek and Jastrzebski, Stanis{\l}aw},
  journal={arXiv preprint arXiv:2002.08264},
  year={2020}
}


@inproceedings{Chen2022sat,
  title={{Structure-aware transformer for graph representation learning}},
  author={Chen, Dexiong and O’Bray, Leslie and Borgwardt, Karsten},
  booktitle={Proceedings of International Conference on Machine Learning},
  pages={3469--3489},
  year={2022},
  organization={PMLR}
}

@article{Gao2023transfoxmol,
    title={{TransFoxMol: predicting molecular property with focused attention}},
    author = {Gao, Jian and Shen, Zheyuan and Xie, Yufeng and Lu, Jialiang and Lu, Yang and Chen, Sikang and Bian, Qingyu and Guo, Yue and Shen, Liteng and Wu, Jian and Zhou, Binbin and Hou, Tingjun and He, Qiaojun and Che, Jinxin and Dong, Xiaowu},
    journal={Brief Bioinform},
    volume={24},
    number={5},
    pages={bbad306},
    year={2023},
}

%%%%%%%%%%%%%%%%%%%%%%%%%%%%%%%%%%%%%%%%%%%%%%%%%%%%%%%%%%%%%%%%%%%%%%%%%


# number={01},
@inproceedings{Zhou2019deep,
  title={{Deep interest evolution network for click-through rate prediction}},
  author={Zhou, Guorui and Mou, Na and Fan, Ying and Pi, Qi and Bian, Weijie and Zhou, Chang and Zhu, Xiaoqiang and Gai, Kun},
  booktitle={Proceedings of the AAAI conference on artificial intelligence},
  volume={33},
  pages={5941--5948},
  year={2019}
}


@article{Wang2022molecular,
  title={{Molecular contrastive learning of representations via graph neural networks}},
  author={Wang, Yuyang and Wang, Jianren and Cao, Zhonglin and Barati Farimani, Amir},
  journal={Nat Mach Intell},
  volume={4},
  number={3},
  pages={279--287},
  year={2022},
  publisher={Nature Publishing Group UK London}
}


@article{bjerrum2017smiles,
  title={{SMILES enumeration as data augmentation for neural network modeling of molecules}},
  author={Bjerrum, Esben Jannik},
  journal={arXiv preprint arXiv:1703.07076},
  year={2017}
}


@inproceedings{Akiba2019optuna,
  title={{Optuna: A next-generation hyperparameter optimization framework}},
  author={Akiba, Takuya and Sano, Shotaro and Yanase, Toshihiko and Ohta, Takeru and Koyama, Masanori},
  booktitle={Proceedings of the 25th ACM SIGKDD international conference on knowledge discovery \& data mining},
  pages={2623--2631},
  year={2019}
}


@article{Vaswani2017attention,
  title={{Attention is all you need}},
  author={Vaswani, Ashish and Shazeer, Noam and Parmar, Niki and Uszkoreit, Jakob and Jones, Llion and Gomez, Aidan N and Kaiser, {\L}ukasz and Polosukhin, Illia},
  journal={Adv Neural Inf Process Syst},
  volume={30},
  year={2017}
}
