## Supplementary material for "CycPeptMP: Enhancing Membrane Permeability Prediction of Cyclic Peptides with Multi-Level Molecular Features and Data Augmentation": latex source: Results_and_Discussion.tex

\section*{Results and Discussion}

\subsection*{Performance comparison}

\begin{table*}[tb]
\caption{Performance comparison between CycPeptMP and seven baseline methods using the test set. The metrics are the averaged values of three repeated runs, and the best result for each metric is indicated in bold.}
\label{tab_results}
\centering
\begin{tabular}{ccccccccc}
\toprule
Evaluation Metrics & RF    & SVM (2D) & SVM (2D3D) & MAT   & SAT   & PharmHGT & FinGAT & CycPeptMP           \\
\midrule
RMSE             & 0.616 & 0.653    & 0.594      & 0.978 & 1.071 & 0.698    & 0.657  & $\mathbf{0.503}$    \\
R                & 0.815 & 0.793    & 0.833      & 0.408 & 0.015 & 0.771    & 0.788  & $\mathbf{0.883}$    \\
\bottomrule
\end{tabular}
\end{table*}


The prediction accuracy results for the test set are shown in Table~\ref{tab_results}, and the prediction accuracy results for the valid set are shown in \SupTabResultsValid.
CycPeptMP ranked first in all evaluation metrics ($\mathrm{RMSE}=0.503$, $\mathrm{R}=0.883$), reflecting a significant improvement in prediction performance over all existing methods.
Considering the structural diversity of the test set, CycPeptMP showed good generalization performance and could learn the complex structures of cyclic peptides which is difficult to apply pre-training through augmentation.
The RF model constructed based on Morgan FP showed good prediction performance and ranked third among all methods ($\mathrm{RMSE}=0.616$, $\mathrm{R}=0.815$). 
SVM with 2D peptide descriptors ($\mathrm{RMSE}=0.653$, $\mathrm{R}=0.793$) had lower prediction accuracy than the RF model, whereas SVM with 3D descriptors improved prediction accuracy, making it superior to the RF model for the test set ($\mathrm{RMSE}=0.594$, $\mathrm{R}=0.833$). 
Cyclic peptide membrane permeation by passive diffusion has a negative correlation with molecule size. 
SVM could predict permeability to some extent by using lipophilicity descriptors, such as LogP, which are largely dependent on molecular weight.
CycPeptMP effectively combined Morgan FP and $16$ 2D and 3D peptide descriptors as peptide-level information to comprehensively characterize peptide structures from both a topological and physicochemical perspective, leading to an improvement in prediction capabilities.
Neither graph representation transformer-based MAT ($\mathrm{RMSE}=0.978$, $\mathrm{R}=0.408$) nor SAT ($\mathrm{RMSE}=1.071$, $\mathrm{R}=0.015$) could predict membrane permeability. 
Although MAT and SAT are state-of-the-art methods for predicting small-molecule properties, they could not effectively learn the structures of more complex cyclic peptides. 
In addition to atom-level information, PharmHGT with fragment MACCS Keys ($\mathrm{RMSE}=0.698$, $\mathrm{R}=0.771$) and FinGAT with molecular Morgan FP ($\mathrm{RMSE}=0.657$, $\mathrm{R}=0.788$) had significantly improved prediction accuracies compared to MAT and SAT. 
FinGAT showed the same level of accuracy as the RF model and 2D SVM.
These results revealed that designing features from various perspectives may be a key to successfully predicting the membrane permeability of cyclic peptides.
These findings indicate that CycPeptMP effectively employs three levels of features to capture a wide range of structural information, from the smallest atomic detail to the broader peptide-level conformation.


\begin{figure}[tb]
    \centering
    \includegraphics[width=0.9\linewidth, natwidth=21.59cm, natheight=17.29cm]{Fig/plot.png}
    \caption{CycPeptMP prediction results for the test set. The predicted value of the test set is the average value of three runs.}
    \label{fig_plot}
\end{figure}


Peptides with an experimental value of $-8$, the lower limit, could not be predicted by CycPeptMP (Figure~\ref{fig_plot}, results for the three validation sets are shown in \SupFigPlotValid).
Compared to the RMSE of all data ($0.503$), the RMSE decreased significantly for data excluding peptides with the experimental value of $-8$ ($0.468$).
Due to the detection limits of the different assay environments, we rounded the data from $-8$ to $-10$ to $-8$, though this may not be the only suitable approach.


\subsection*{Ablation study of atom and monomer models}

\begin{figure}[tb]
    \centering
    \includegraphics[width=0.9\linewidth, natwidth=19.44cm, natheight=15.24cm]{Fig/ablation-atom&monomer.png}
    \caption{Ablation results (RMSE) for the atom and monomer models using the test set.}
    \label{fig_ablation-atom&monomer}
\end{figure}


We conducted ablation studies on atom and monomer models with complex architectures (Figure~\ref{fig_ablation-atom&monomer}, the results for the validation set are shown in \SupFigAblationValid~(A)). 
For the atom model, A is the original model and A--aug is the result without data augmentation. 
We measured the prediction accuracy when not using the $Bond$ matrix (A--bond), using ordinary absolute positional encoding \cite{Vaswani2017attention} instead of the $Bond$ matrix (A--abpe), retaining only the $Conf$ block (A--graph), or retaining only the graph block (A--conf).
As shown in Figure~\ref{fig_ablation-atom&monomer}, the prediction accuracy of the atom model was significantly improved by augmentation (A: $0.653$, A--aug: $0.931$).
Regarding the architecture changes of the atom model, the original A showed the highest prediction accuracy, and any element deletion decreased the prediction accuracy.
It was better to capture the relationship between atoms using $Bond$ (A: $0.653$) than absolute positional encoding (A--abpe: $0.678$), and the impact of removing $Graph$ block (A--graph: $0.67$) was greater than that of removing $Conf$ block (A--conf: $0.659$).


For the monomer model, M is the original model and M--aug is the result without data augmentation. 
We also measured the change in accuracy when replacing the general 1D-CNN layers with CyclicConv layers (M--Cyclic). 
Similar to the atom model results, the prediction accuracy of the monomer model was significantly improved by augmentation (M: $0.57$, M--aug: $0.891$). 
These results showed that SMILES enumeration for the atom model and sequence arrangement for the monomer model effectively improved learning efficiency, and that the augmentation technique is essential for learning the complex structure of cyclic peptides. 
Additionally, the 1D-CNN layer (M: $0.57$) was superior to the CyclicConv layer (M--Cyclic: $0.632$), consistent with previous findings \cite{Li2022}.


\subsection*{Ablation study of fusion model}


\begin{figure}[t]
    \centering
    \includegraphics[width=0.9\linewidth, natwidth=10.72cm, natheight=15.06cm]{Fig/ablation-fusion.png}
    \caption{Ablation results (RMSE) for the fusion model using the test set. (A) Results for different numbers of input replicas. (B) Results for different architectures.}
    \label{fig_ablation-fusion}
\end{figure}


The ablation study for the fusion model measured the influences of the number of replicas generated by augmentation and changes in architecture.
Figure~\ref{fig_ablation-fusion}~(A) shows the accuracy based on $1$ (no augmentation), $5$, $10$, $20$, $30$, $40$, $50$, and $60$ (CycPeptMP) replicas per peptide. 
Even with five replicas (F--5: $0.544$), we observed a significant improvement in prediction accuracy compared to that without augmentation (F--1: $0.61$).
However, over $20$ replicas showed approximately the same prediction accuracy as the amount of training data increased, which may be due to the limitations of increasing diversity by mere reordering of inputs and lack of diversity in the generated conformations.


In Figure~\ref{fig_ablation-fusion}~(B), F is the original CycPeptMP model; F--aux is that without auxiliary loss; F--atom, F--mono, and F--pep are those lacking the respective sub-models; F--3D is that not using all 3D information ($Conf$ and 3D descriptors); and F--ensem is that where each sub-model is allowed to directly predict membrane permeability and the average ensemble of three predictions is taken.
% validだと上がった
Auxiliary loss did not improve prediction accuracy (F--aux: $0.501$).
Furthermore, the prediction accuracy decreased when removing any of the three sub-models, indicating that the three levels of information is essential for predicting membrane permeability. 
The peptide model had the greatest influence (F--pep: $0.539$), followed by the monomer (F--mono: $0.53$) and atom model (F--atom: $0.511$).
% The results of the RF ($0.616$) and SVM models ($0.594$) (Table~\ref{tab_results}) and the results of the atom (A: $0.653$) and monomer models (M: $0.57$) (Figure~\ref{fig_ablation-atom&monomer}) collectively emphasize the difficulty in predicting the permeability of cyclic peptides using only atom graphs, similar to that in predicting small-molecule properties.
The use of 3D information improved prediction accuracy (F--3D: $0.528$), though not significantly so.
Accuracy may be improved by generating conformations using a more rigorous method, such as MD simulations.
Finally, the average prediction accuracy decreased further when using a sub-model ensemble (F--ensem: $0.529$).
This indicates that, rather than having each sub-model predict permeability directly, it was better to extract latent features.


\subsection*{Comparison with MD-based method}

Cyclic peptides tend to exist in various conformations, resulting in slow conformational transitions relative to simulation time scales. 
Sugita et al., in the first large-scale prediction of cyclic peptide membrane permeability \cite{Sugita2021}, used steered MD and replica-exchange umbrella sampling to accelerate sampling and simulated the membrane permeation process of $100$ six-residue and $56$ eight-residue peptides through a lipid bilayer. 
We compared their prediction results with the CycPeptMP results for $23$ peptides (\SupTabResultsMD) included in three validation sets ($16$ peptides) and the test set ($7$ peptides). 


While the MD-based method could not predict the membrane permeability of these $23$ peptides ($\mathrm{RMSE}=1.899$), CycPeptMP was able to make accurate predictions for all of them ($\mathrm{RMSE}=0.127$) (Figure~\ref{fig_MD}).
Because hydrophobic cyclic peptides have insufficient solubility, diffuse slowly in the unstirred water layer, and are likely adsorbed to the membrane, Sugita et al. could not reproduce these behaviors using the inhomogeneous solubility-diffusion model (ISDM), which considers only direct membrane permeation processes \cite{Sugita2021}.
Thus, they reported a prediction accuracy ($\mathrm{R}$) of only $0.21$ for all $100$ six-residue peptides; however, when $33$ hydrophobic peptides ($\mathrm{AlogP}>=4$) were excluded, the accuracy increased to $0.54$. 
A similar trend was seen among the $23$ peptides compared in this study: by excluding six peptides with $\mathrm{AlogP}>=4$, the $\mathrm{RMSE}$ of the MD-based method increased from $1.899$ to $1.219$.
Overall, CycPeptMP could accurately and quickly predict peptide permeability with far superior performance compared to MD-based methods, and thus represents a promising tool for cyclic peptide drug discovery.


\begin{figure}[tb]
    \centering
    \includegraphics[width=0.8\linewidth, natwidth=11.25cm, natheight=18.3cm]{Fig/MD.png}
    \caption{Prediction results of (A) MD-based method and (B) CycPeptMP.}
    \label{fig_MD}
\end{figure}


% \subsection*{Case study}
