## Supplementary figures and images for "CycPeptMP: Enhancing Membrane Permeability Prediction of Cyclic Peptides with Multi-Level Molecular Features and Data Augmentation"

### ablation-atom&monomer.png

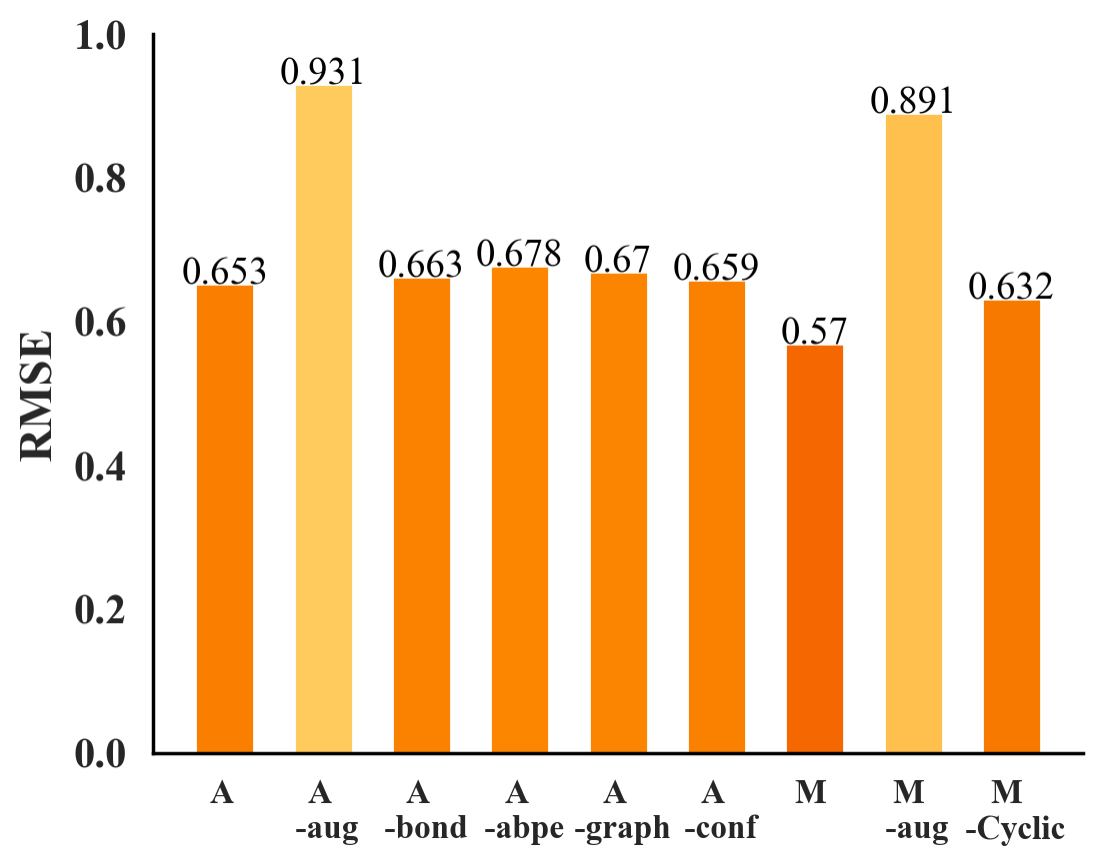

### ablation-fusion.png

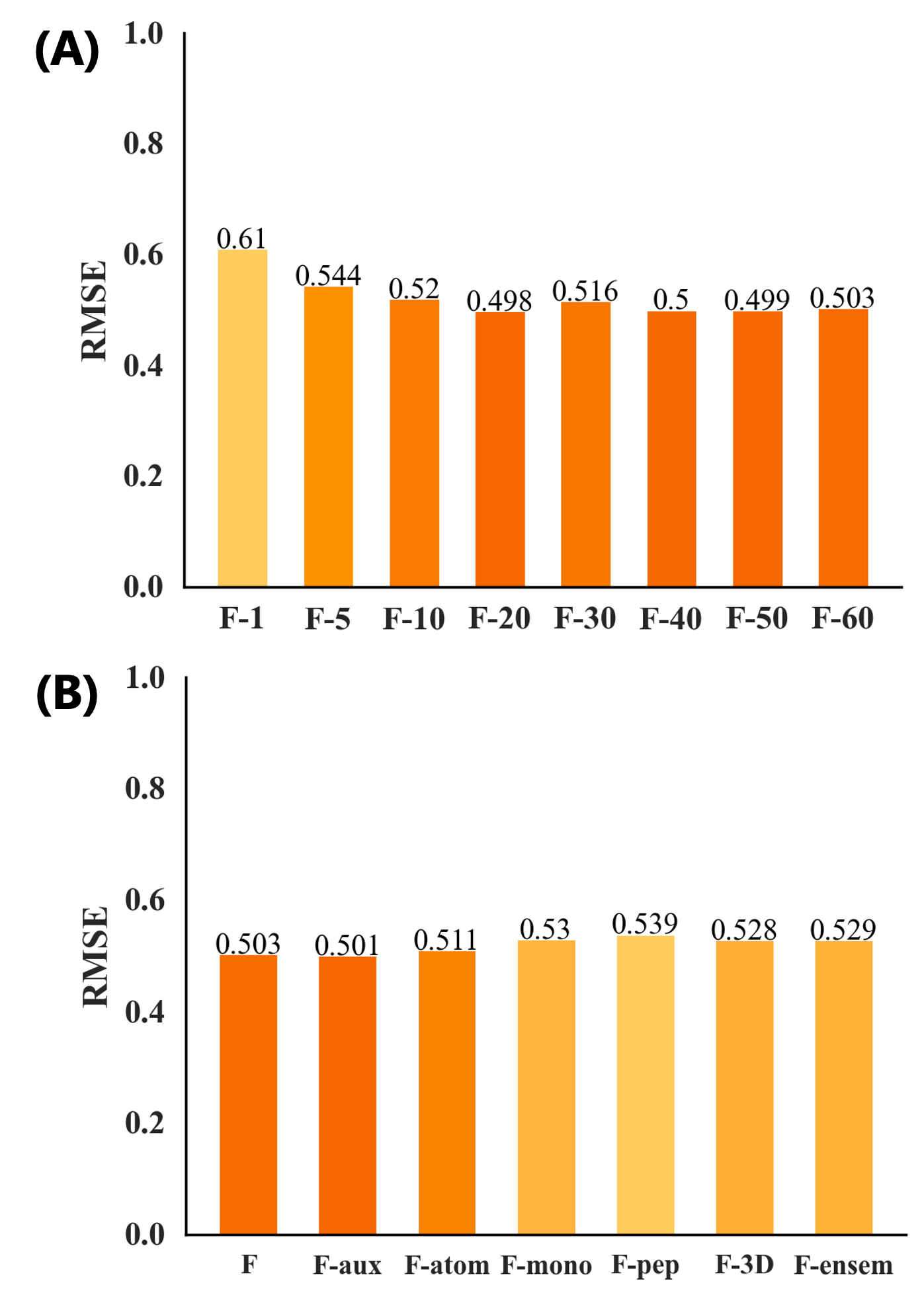

### augmentation.png

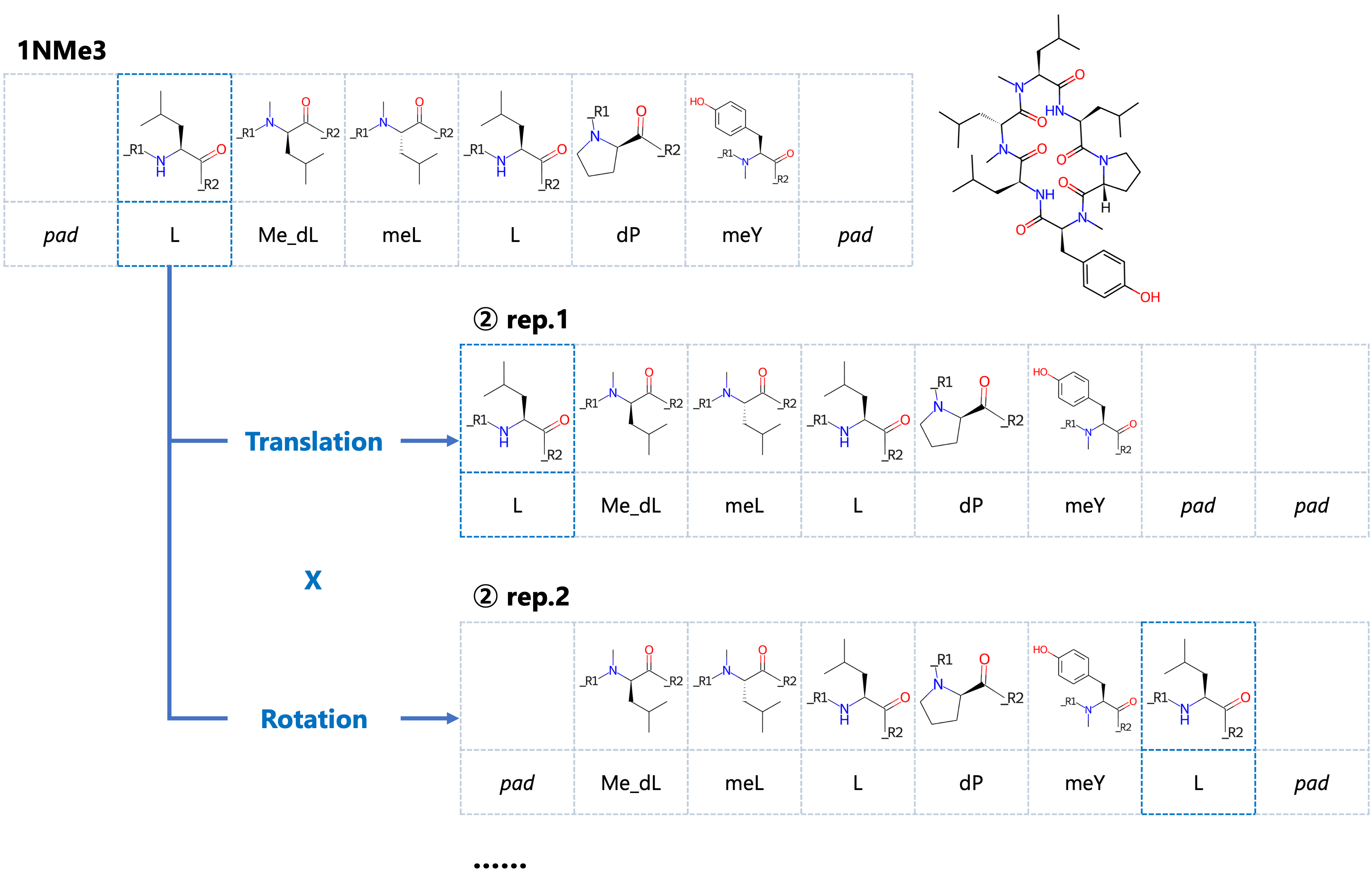

### framework.png

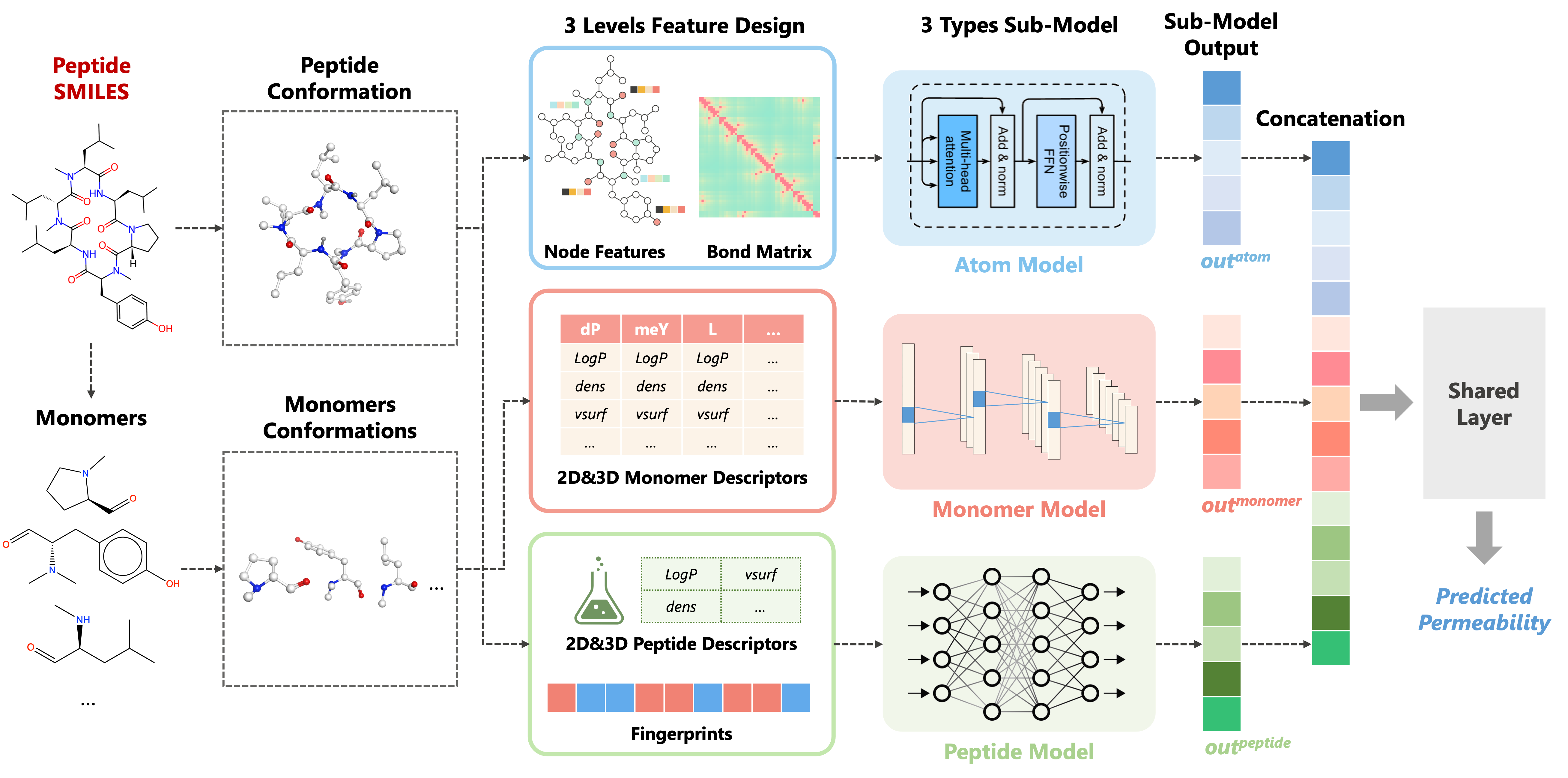

### MD.png

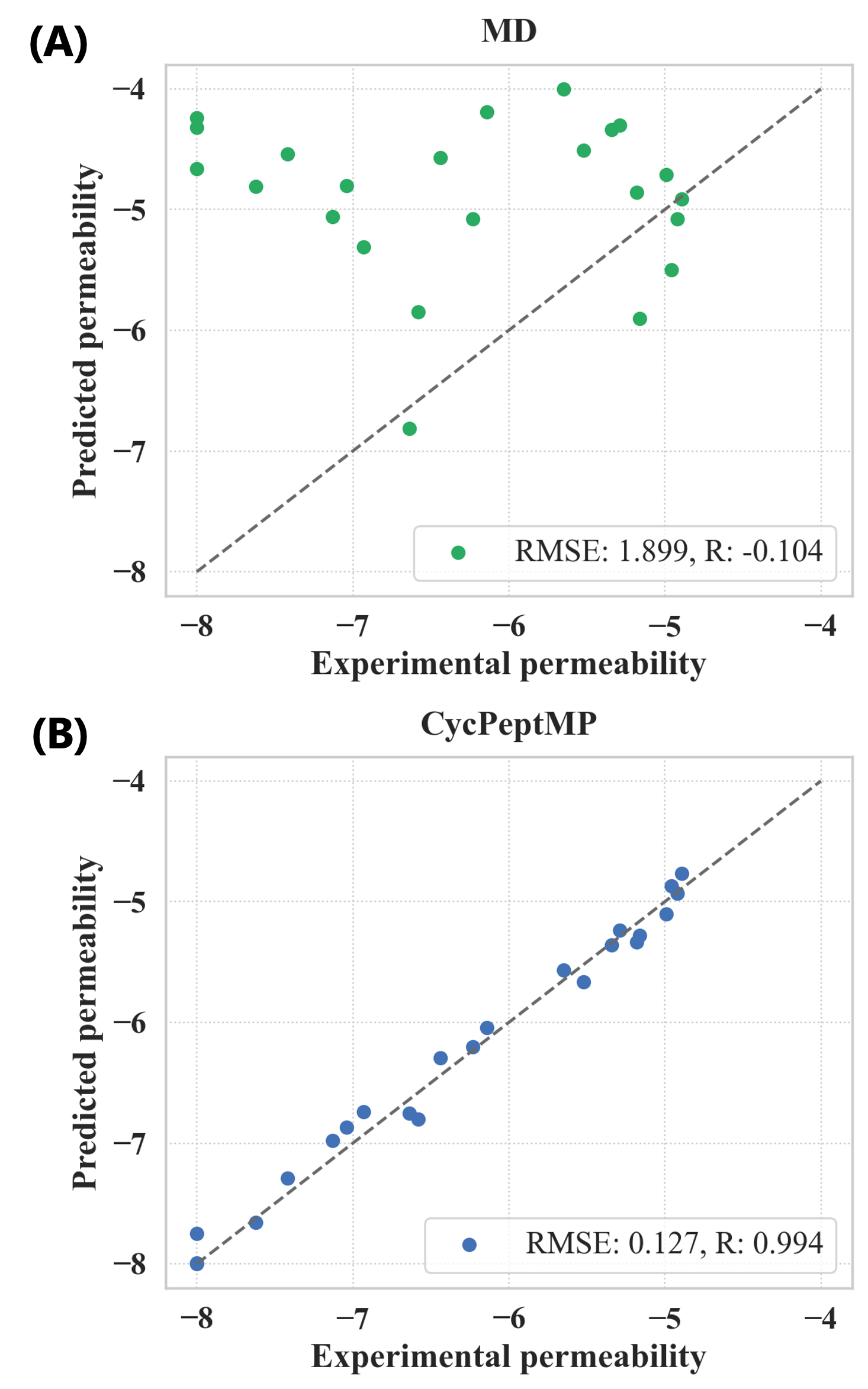

### model.png

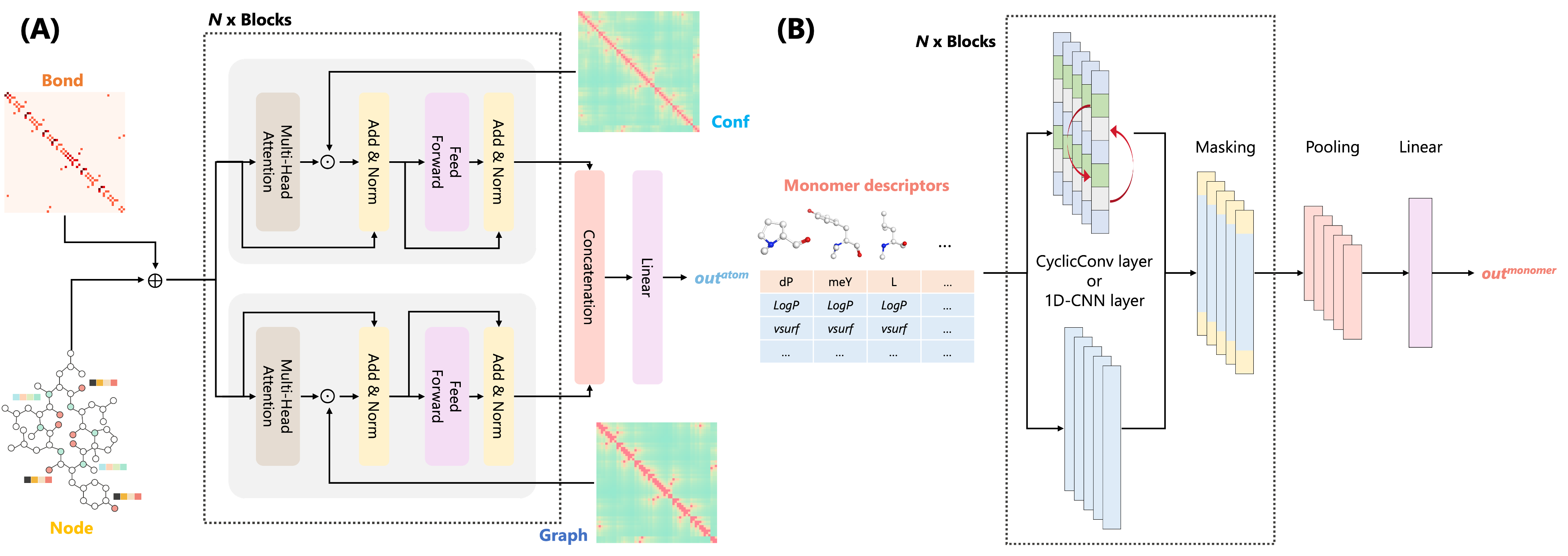

### plot.png

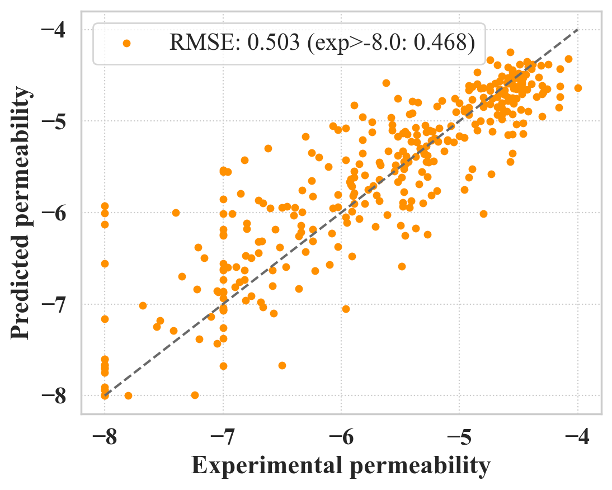

### set-distribution.png

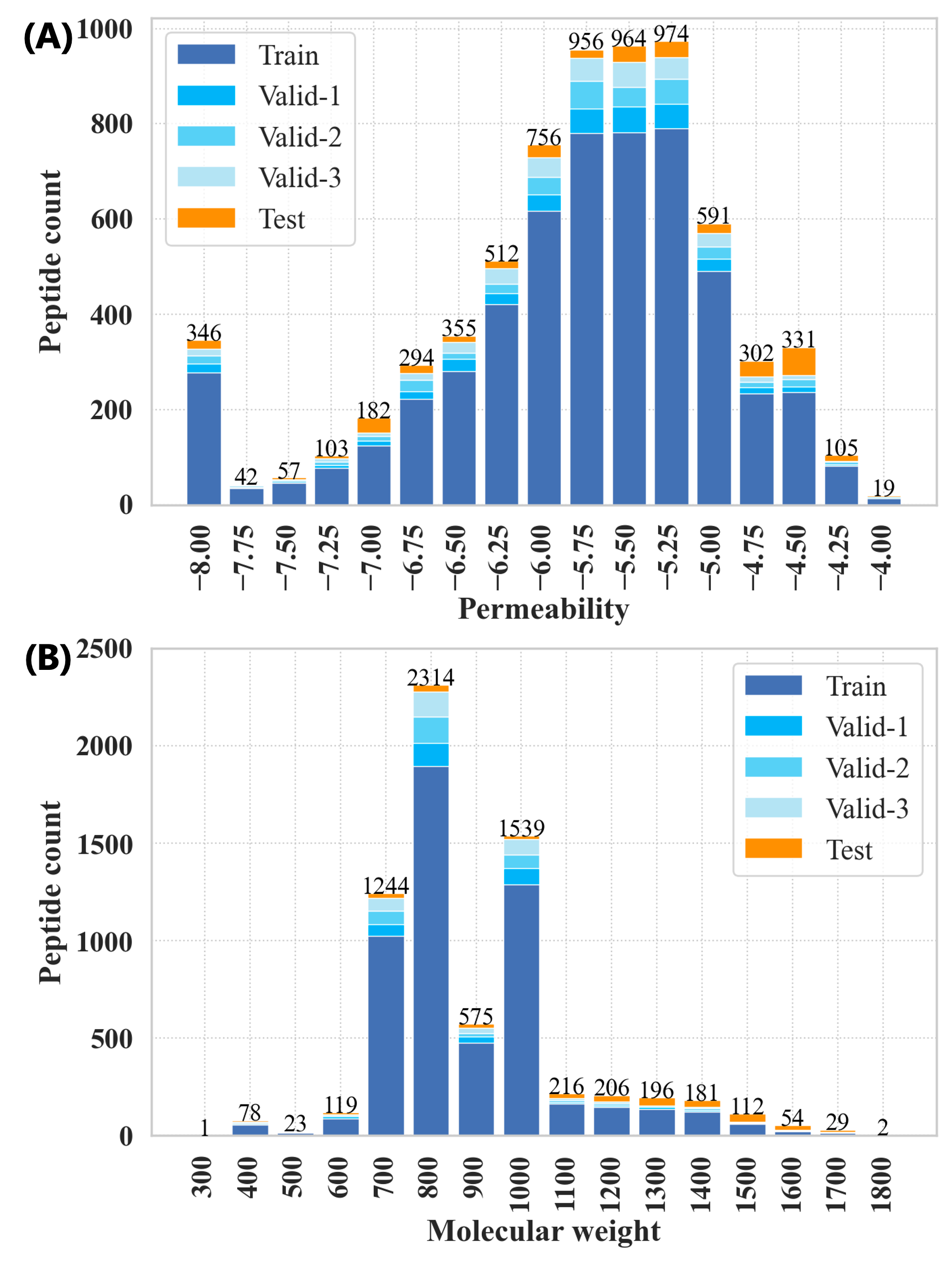
